# A connectomics-driven analysis reveals novel characterization of border regions in mouse visual cortex

**DOI:** 10.1101/2024.05.24.595837

**Authors:** Neehal Tumma, Linghao Kong, Shashata Sawmya, Tony T. Wang, Nir Shavit

## Abstract

Leveraging retinotopic maps to parcellate the visual cortex into its respective sub-regions has long been a canonical approach to characterizing the functional organization of visual areas in the mouse brain. However, with the advent of extensive connectomics datasets like MICrONS, we can now perform more granular analyses to better characterize the structure and function of the visual cortex. In this work, we propose a statistical framework for analyzing the MICrONS dataset, particularly the V1, RL, and AL visual areas. In addition to identifying several structural and functional differences between these regions, we focus on the *borders* between these regions. By comparing the V1-RL and RL-AL border regions, we show that different boundaries between visual regions are distinct in their structure and function. Additionally, we find that the V1-RL border region has greater synaptic connectivity and more synchronous neural activity than the V1 and RL regions individually. We further analyze structure and function in tandem by measuring information flow along synapses, observing that the V1-RL border appears to act as a bridge between the V1 and RL visual areas. Overall, we identify numerous measures that distinguish the V1-RL border from the larger V1-RL network, potentially motivating its characterization as a distinct region in the mouse visual cortex.

## 1. Introduction

The mammalian visual cortex is among the most advanced visual processing systems in existence. It comprises multiple visual areas, including the primary visual cortex (V1), which directly receives most of the visual inputs from the thalamus, and other higher order visual areas. In mice, such higher order visual areas include the rostrolateral (RL), anterolateral (AL), and lateromedial areas (Marshel et al., 2011). These regions have highly complex synaptic connections within and between each other (Nassi and Callaway, 2006) and display unique functional characteristics, with one of the most notable being retinotopy (Tyler, 2004).

Retinotopy is a property in which portions of the visual field are each represented by and encoded within specific areas in the visual cortex (Fize et al., 2003). Classically, retinotopy is most often determined through experiments including the rotating bar and expanding ring stimuli (Tyler et al., 2005), though more recent work allows retinotopic mappings to be recovered from natural movie stimuli (Kumar et al., 2021). Retinotopic maps are canonically used to induce a parcellation of the visual cortex into its sub-regions. Such maps allow for the characterization of the spatial arrangement of neurons in the visual cortex, of defining boundaries between different visual areas, and of determining what type of stimuli different neurons respond to (Marshel et al., 2011; Garrett et al., 2014). Similarly, architectonics is a field that utilizes structural information to discern different visual regions. Most techniques utilize the staining of different neurofilaments and receptors, the tracing of broad cortical connections, and the quantification of neuronal density in different regions (Baldauf, 2005; Van der Gucht et al., 2007; Wang et al., 2007; Herculano-Houzel et al., 2013; Zilles and Palomero-Gallagher, 2017). However, recent work has shown that there is significant variation in the retinotopically discerned regions across different animals in anatomical space (Garrett et al., 2014), and even disagreements between architectonically and retintopically discerned boundaries within the same animal (Zhuang et al., 2017).

To better explore the intersection of architectonics and retinotopy—of function and structure—in this work, we utilize a more recently developed framework for analyzing the visual cortex (and more generally the brain) known as *connectomics*. In particular, we strive to understand how connectomics can be used to improve upon our existing characterization of the mouse visual cortex currently enabled by retinotopy and architectonics, especially where they disagree. Connectomics is an emerging field at the intersection of neuroscience and computer science that leverages high resolution electron microscopy to characterize exact neuron-to-neuron connections (Behrens and Sporns, 2012). One of the most comprehensive connectomics datasets is from the Machine Intelligence from Cortical Networks (MICrONS) consortium. It comprises a cubic millimeter of the mouse visual cortex and includes over 75,000 neurons and 500,000 synapses. Furthermore, it is one of the few connectomics datasets to also include functional traces associated with individual neurons (Consortium et al., 2021). Recent work has utilized the MICrONS dataset to investigate both general wiring rules (Ding et al., 2024) and those of inhibitory circuits in particular (Schneider-Mizell et al., 2023).

Connectomics differs from retinotopy in that it goes beyond studying the functional organization of specific sensory regions by providing a comprehensive and highly detailed view of the brain’s structural and functional connectivity patterns (Dorkenwald et al., 2023). While retinotopy focuses on how sensory information is represented in specific brain areas, connectomics offers an unprecedented level of detail that includes connectivity data in addition to functional data (Scholtens and van den Heuvel, 2018). Connectomics maps the intricate network of individual neurons, their synapses, and the broader patterns of connectivity, surpassing the resolution of functional imaging methods like retinotopy. This more granular approach allows us to explore the fine-scale architecture of neural circuits, revealing the precise organization of connections and offering a holistic understanding of how information flows and is processed throughout the brain.

In this work, we use the unique opportunity presented by the MICrONS dataset to profile the structural and functional granularities of the mouse visual cortex and use this information to understand if regions in the mouse visual cortex delineated via retinotopy exhibit structural and functional differences when analyzed under the lens of connectomics. We place our attention on the V1, RL and AL visual areas and investigate the biological neural network induced by the portion of the connectomics data corresponding to these retinotopically-induced regions. In doing so, we focus much of our analysis on the borders between these regions: although retinotopic boundaries between visual regions have been shown to demonstrate unique phenomena such as mirroring and compression (Lyon et al., 2002), they are otherwise little understood. Furthermore, discrepancies between retinotopic maps of different members of the same species and between retinotopic and architectonic maps for the same animal are most apparent at the boundaries (Garrett et al., 2014; Zhuang et al., 2017). Using MICrONS, we draw novel conclusions about anatomical and functional patterns present at the V1-RL and RL-AL borders that have yet to be properly characterized previously.

## 2. Main Findings

In this section, we provide a brief overview of the main findings in this paper before discussing them in further detail. We conduct a novel combined analysis of the structural and functional components of the mouse visual cortex using MICrONS, placing our attention on the V1, RL, and AL visual areas, as well as the boundaries between them. We find that:

1. The V1-RL boundary possesses heightened synaptic connectivity and more pairwise correlated neural activity than the rest of the V1-RL network. In contrast, we observe neither of these trends at the RL-AL border (Section 4.1).
2. The V1 and RL regions are distinct with respect to a variety of features, including neuron and synaptic density, degree of functional correlation, cell type distributions, information propagation rate, and the distance between neighboring neurons (Section 4.2).
3. The V1-RL border presents itself uniquely with respect to both structure and function, a notion that is not captured via parcellations over retinotopic or architectonic maps (4.3, Appendix E.2). Additionally, we find that this characterization is robust—it is identifiable among fake border regions constructed via translation and rotation (Section 4.5, Appendix D)
4. The V1-RL border region appears to act a bridge for communication between the V1 and RL visual areas. We find that the information flow along synapses traversing the border is the highest amongst all synapses in the V1-RL network (Section 4.4).

These findings provide a statistical characterization of the V1, RL, and AL visual regions to a degree that was previously not possible due to a lack of connectomics data like MICrONS.

## 3. Methods

### 3.1. Dataset description and construction

The connectomics dataset that we utilize is from the Machine Intelligence from Cortical Networks (MICrONS) consortium. It includes more than 75,000 neurons and 500,000 synapses. It also includes functional data in the form of calcium and derived activity traces for all neurons when the mouse was viewing natural movie stimuli. However, only approximately 33,000 neurons have their functional and anatomical information paired together. These neurons are considered co-registered. Each neuron has its position recorded in 3D anatomical space, as well as a list of synaptic connections it has with its neighbors. See Section Appendix A for a list of neuron cell types within MICrONS. MICrONS uses a convention such that *x* and *z* coordinates correspond to a point on the flattened cortical surface, and the *y* coordinate indicates cortical depth. The connectome was reconstructed from high-resolution electron microscopy (EM) images. To distinguish different visual areas (i.e. V1, RL, AL) via retinotopy, the MICrONS group used a drifting bar stimulus to generate a sign map, and then manually annotated traces within that map.

For the purposes of our analysis, we will visualize the dataset through a cortical surface view (i.e. the *x* and *z* dimensions) and aggregate along cortical depth (i.e. the *y* dimension; see Figure 1). This is because we find that the meaningful variations for our analyses exist primarily in the directions along the cortical sheet. However, note that we perform some analyses on a per-layer basis, which will be clarified when necessary. Finally, since cortex depth is not perfectly aligned with the y-axis in MICrONS, we rotate the volume about the border between V1 and RL such that the cortical layers are aligned with the *xz* plane.

**Figure 1:**
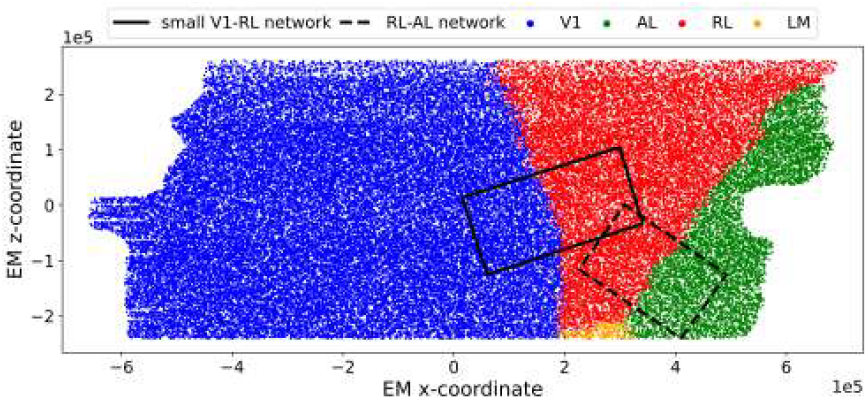
Visualization of the entire MICrONS dataset in EM-space, labeled by visual area (the axes are in units of *µm*). This is a flattened view of the cortical surface. The V1-RL and RL-AL networks/windows denote the portion of the dataset we analyze in the first part of this work.

In the first part of our work, we restrict the scope of our analysis to the portion of the V1-RL and RL-AL networks shown in Figure 1. We orient the regions of interest such that the borders are roughly parallel to the left and right sides of their respective windows, and are roughly equal in size. We do so to ensure that analyses comparing the two regions are not confounded by factors such as the orientation of the border or differences in area.

In the analysis comparing the V1-RL and RL-AL networks, the size of the windows are bottlenecked by the data availability of the AL visual area within MICrONS (Figure 1). For a more in-depth analysis, we restrict the scope of our analysis to the V1-RL network and capture as much of the dataset as possible. However, we also need to take into account edge effects, where neurons near the edges of the dataset have fewer connections due to many of their neighboring neurons not being present in MICrONS. Furthermore, we note that there is a region within the dataset with a high degree of manual proofreading that resides primarily in the center of the V1 region. The V1-RL window, shown in Figure 2, is constructed to capture as much data as possible while minimizing any potential confounding factors, such as edge effects and a difference in manual proofreading.

**Figure 2:**
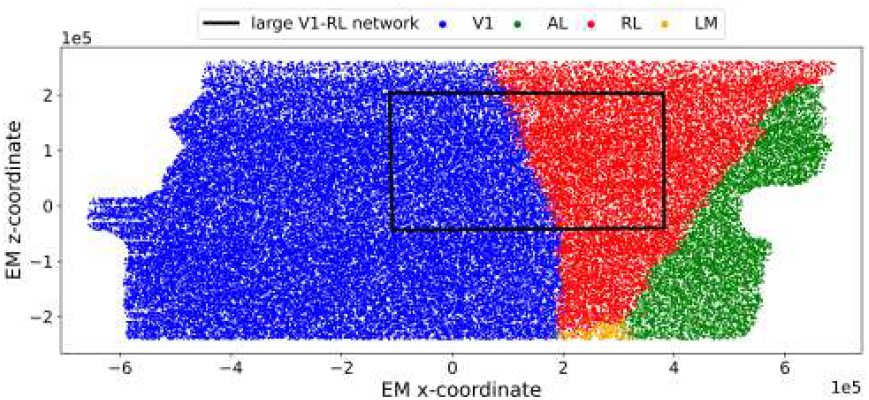
The portion of the network we analyze when focusing solely on the V1-RL network (as compared to Figure 1).

### 3.2. Fitting the borders

In the cortex surface view, the retinotopic area assignments for each neuron in the windows we consider (Figure 1, 2) include a boundary that can be approximated quite closely by a linear function of the *x* and *z* coordinates. Using a linear function allows us to perform simple translations and rotations on the border. We fit the borders as a binary classification problem on the area assignment for every neuron, not only the coregistered ones. If we let *w* denote the vector of V1/RL area labels, *x* denote the vector of EM *x* coordinates and *z* denote the vector of EM *z* coordinates, the functional form of the border is given by a logistic regression as follows:

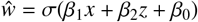

### 3.3. Border regions as “cuts” and “windows”

Throughout this paper, we investigate two possible interpretations of the borders between visual regions—that of a “cut” and that of a “window”. Previous works using retinotopic data have framed the characterization of the mouse visual cortex and its sub-regions as a parcellation problem of individual neurons to different regions (Garrett et al., 2014; Kumar et al., 2021). In this context, there are two reasonable interpretations for the V1-RL and RL-AL borders: the cut interpretation (readily derived from a retinotopic parcellation) and the window interpretation (which less clearly follows from a retinotopic parcellation).

Under the cut interpretation, a border serves as a divider between two retinotopically distinct regions. In other words, it effectively acts as a decision boundary between the two regions. The cut in this instance refers to the plane fit using visual area labels, which were collected from a parcellation computed on retinotopic data (Figure A8A). Recall the fitting procedure for the border from Section 3.2.

Under the window interpretation, the volume surrounding the border plane is now assigned significant importance. We now consider all neurons within distance *d* of the plane as members of a border window (Figure A8E). *d* refers to the shortest Euclidean distance between a point and the plane of the fit border. This formulation now assigns importance to a volume of area that we can consider as a separate region, distinct from the two regions the border delineates. We go into more detail about these two interpretations in Appendix E.

### 3.4. Metric definitions

Here, we define various structural and functional metrics we use in our analyses.

#### 3.4.1. Graph representation of dataset

Consider a graph representation of the MICrONS dataset. Due to the inclusion of synaptic data, we can consider MICrONS as a directed graph, *G* = {*V, E*}. *V* is the set of all neurons in the dataset, and *E* is the set of all edges between neurons. For any neurons *v*_1_, *v*_2_ ∈ *V*, there exists an *e*_12_ ∈ *E* if and only if there is a synapse going from *v*_1_ to *v*_2_. In this context, we refer to *v*_1_ as the *pre-synaptic* neuron and *v*_2_ as the *post-synaptic* neuron. We will use vertex and neuron interchangeably, as well as edge and synapse.

The graph *G* represents the full dataset. Recall that only a subset of it is coregistered; so, we denote this subgraph by *G*_*F*_ = {*V*_*F*_, *E*_*F*_} ⊆ *G*. Since the coregistered subgraph has functional data, we can represent a neuron *n* ∈ *V*_*F*_ as a tuple given by (*v, t*) where *v* is the vertex in the graph corresponding to the neuron and *t* is the trace of functional activity corresponding to the neuron’s activations induced by natural movie stimuli. We preprocess the activity traces by setting all measurements less than 10^−3^ to 0, as we consider measurements below this threshold as noise during recording.

We compute estimators for the metrics detailed below using spatial bootstrapping on the graph (Thompson et al., 2016). This entails sampling sub-volumes of the 3D graph that represents the network. In particular, if we denote the dimensions of the subgraph volume by (*x, y, z*), then a randomly sampled sub-volume’s dimensions are given by (*tx, y, tz*) for hyperparameter *t* such that 0 *< t <* 1 (note that *x, y, z* here refer to the dimensions defined in Section 3.1). Bootstrapping in this manner captures spatial correlations present in the data, enabling us to construct viable tests of statistical significance.

#### 3.4.2. Neuron Density

The neuronal density of a region *R* with *N* total neurons and a volume of *V* is calculated as:

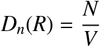

#### 3.4.3. Synaptic density

The synaptic density of a directed graph *G* with *V* vertices and *E* edges is calculated as:

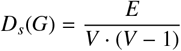

#### 3.4.4. 2-cycle density

The 2-cycle density (*C*_2_) of a directed graph *G* is calculated as follows:

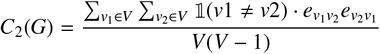

where *V* is the set of vertices in the graph and *e*_*ij*_ is 1 if and only if there is an edge going from node *i* to node *j* and 0 otherwise.

#### 3.4.5. Distribution over 3-motifs

Let *G* be a network with *N* nodes. A 3-motif is a subgraph *M* of *G* consisting of three nodes and the edges between them (Figure A1). The distribution *P*(*M*) over 3-motifs in the network *G* can be calculated by counting the occurrences of each distinct 3-motif *M* and normalizing by the total number of possible subgraphs of size 3.

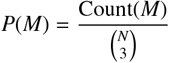

#### 3.4.6. Microcircuit expansions

Let the set of neuron types be denoted by *T* = {2*/*3, 4, 5*/*6, inh} where here we have grouped L5 and L6 neurons together. In this work, we will consider the following two microcircuits:

- 4 → 5/6 → 2/3 → 4
- 2/3 → inh → 4 → 2/3

Both of these circuits are theorized units of visual information processing (Hooks and Chen, 2020), with the former a general visual processing circuit and the latter a predictive coding circuit (Bastos et al., 2012). We can more generally represent these circuits as *A* → *B* → *C* → *A* for neuron types *A, B, C*. We define the expansion rate of such as a circuit as follows:

Consider a decomposition of *G* into sub-graphs *G*_*A*_, *G*_*B*_, *G*_*C*_, where each *G*_*L*_ corresponds to a graph comprised only of neurons of type *L*. Select an arbitrary neuron from *G*_*A*_ as the starting point and perform a breadth-first search (BFS) from this node. The BFS is structured to expand in a fixed order: first, it visits neurons in *G*_*B*_; next, it explores neurons in *G*_*C*_; and finally, it reaches additional neurons in *G*_*A*_. The number of neurons encountered at each stage of the BFS is summarized by the 4-tuple

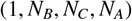

where 1 represents the starting neuron in *G*_*A*_; *N*_*B*_ is the number of *G*_*B*_ neurons reached in the first level of expansion; *N*_*C*_ is the number of *G*_*C*_ neurons reached in the second level of expansion; and *N*_*A*_ is the number of *G*_*A*_ neurons encountered in the third level of expansion. This 4-tuple forms the basis for computing the expansion rate of the circuit for the single neuron *v* as follows:

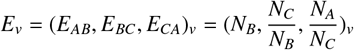

To compute the expansion rate of this circuit over the entire *G* = {*V, E*}, we element-wise average over all possible starting nodes:

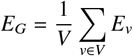

The expansion rate of these microcircuits provides us with estimates of the rate of signal propagation in units that are relevant to visual information processing.

#### 3.4.7. Activity level

Recall that a neuron *n* ∈ *V*_*F*_ has a corresponding trace *t* representing its functional activity. *t* ∈ ℝ^*T*+^ where *T* = 40, 000, representing how many frames there are in the functional data. We can define the average activity *A*(*G*_*F*_) in the graph as follows

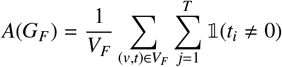

#### 3.4.8. Average pairwise correlation

The pairwise correlation coefficient *r* between two neural traces *X* and *Y* can be calculated using the Pearson correlation formula:

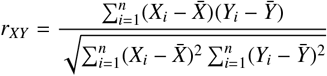

We are interested in the average pairwise correlation over an entire graph *G*_*F*_. Letting *N* = *V*_*F*_, this is computed as

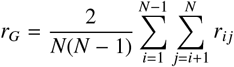

#### 3.4.9. Median information flow (Granger causality)

Pairwise correlation only considers data points between two variables individually, and does not consider temporal influences from one variable to the other. To potentially better capture this, we use Granger causality as a more straightforward measure for the strength of directed information flow (Seth et al., 2015). Consider a pair of neural traces *X* and *Y* corresponding to neurons *n*_*x*_ and *n*_*y*_ respectively such that there exists an edge going from *n*_*x*_ to *n*_*y*_. We let *g*_*XY*_ denote the p-value of a Granger causality test from *X* to *Y* computed for a fixed lag 𝓁. Here we use 𝓁 = 5, which aligns with previous works such as Chen et al. (2023). The median pairwise Granger causality measures the median directional influence of one neuron on another. We compute the median as opposed to the average, due to the skewness of the empirical distribution over p-values we observe. If we let [*g*_*i j*_] denote a list of all pairwise Granger causalities, then the median information flow is computed by

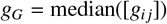

Note that lower values under this metric indicate higher Granger causality and stronger information flow.

#### 3.4.10. Effective dimensionality of neural activity

The effective dimensionality of a set of neural traces (Jazayeri and Ostojic, 2021) is the cumulative explained variance of the top *k* principal components of the trace data for a fixed *k*. This is given by

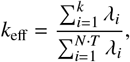

where *N* is the number of traces in the dataset, *T* is the number of time points in each trace, *k* is the number of principal components being considered, and *λ*_*i*_ represents the eigenvalue of the *i*-th principal component, sorted in descending order. Note that higher values of this metric indicate lower dimensional activity.

### 3.5. Sliding window framework

To characterize the the potentially more granular changes in structure and function that may be present in the network that is closer to the border, we construct a sliding window framework.

Using the fitted border (Section 3.2), we can create a window around the border plane where all points within distance 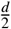 are included, which yields a window of width *d* (Figure 3). Let us denote this window by *W*_*d*_(0). We can generalize this to an arbitrary window over the volume denoted by *W*_*d*_(*T*) where *T* denotes a *T*-unit horizontal translation from the border window *W*_*d*_(0).

**Figure 3:**
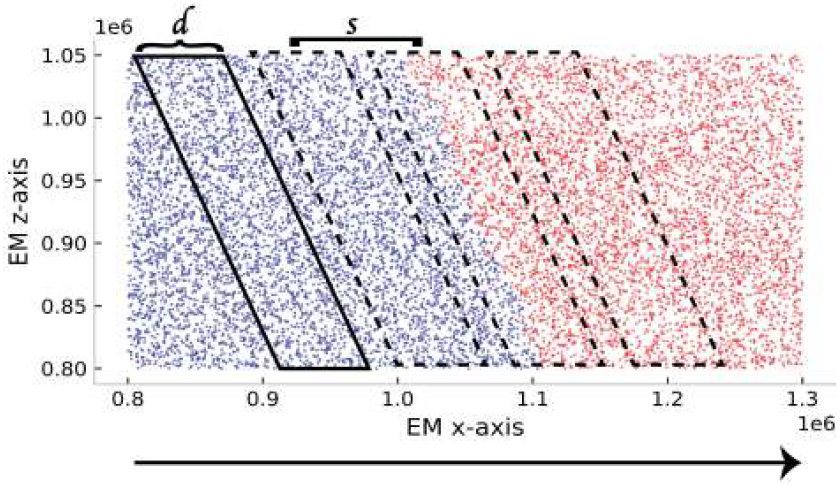
Depiction of sliding window used for analysis. Window is parameterized by width *d* and stride *s*.

Let the leftmost EM x-coordinate in the dataset be *L* and the rightmost be *R*. Denote the x-coordinate corresponding to the border by *B*. We define a sliding window on the V1-RL network as a set of *n* + 1 windows given by

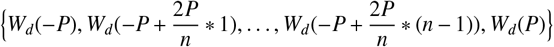

where *P* = min(|*B* − *R*|, |*B* − *L*|). Now, what we have is a set of *n* equally spaced windows that span the network. For any given window *W*_*d*_(*T*), we will refer to the plane that divides it by *C*_*d*_(*T*). This means that we can denote the border plane fit in Section 3.2 by *C*_*d*_(0).

We can compute our metric of choice in each window to create a fine-grained characterization of the evolution of structure and function across the network. Also, note that if we choose an odd number of windows, *W*_*d*_(0) is included in the sliding window set.

### 3.6. Border identifiability experiments

#### 3.6.1. Construction of fake borders

A persistent issue in constructing borders between visual areas is that there is a certain degree of arbitrariness. Previous works have shown that the border fit using anatomical data does not necessarily align with the border fit using retinotopic data (Zhuang et al., 2017). To demonstrate the robustness of our connectomics-driven characterization of the border, we construct false boundaries to see if any of them can be construed as the true border. We denote a fake border cut as *F*_*C*_(*T, θ*) and a fake border window as *F*_*W*_ (*T, θ*) in which our parameterization consists of a horizontal translation *T* from the true border and an angle *θ* which denotes the rotation from the true border (Figure 4A, Figure 4B).

**Figure 4:**
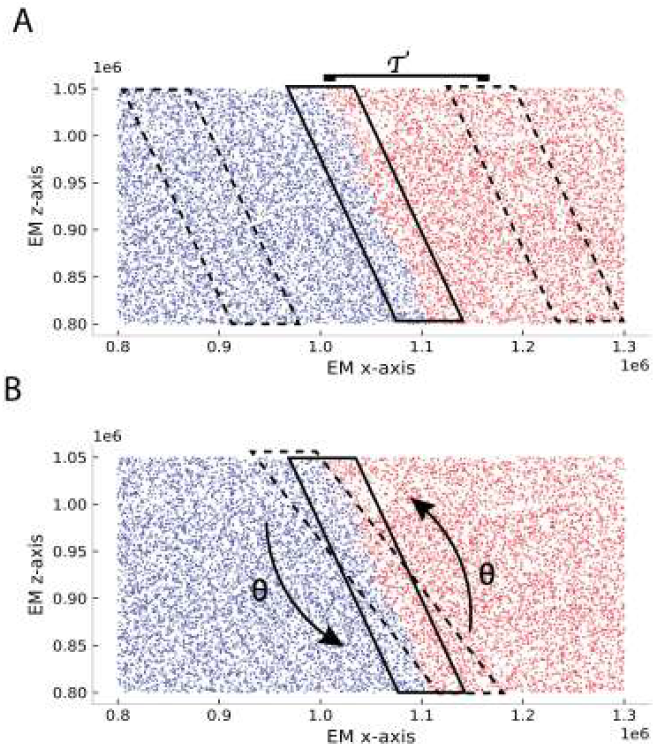
(A) Fake border parameterized by horizontal translation *T*. (B) Fake border parameterized by rotation angle *θ*.

#### 3.6.2. Fake border classifiers

Here, we describe our methodology for strengthening our characterization of the V1-RL boundary. We construct predictive models to attempt to discriminate the true boundary from perturbations constructed via translation and rotation to demonstrate the robustness of our characterization of the true border under the window interpretation as described in Section 3.3. In particular, consider a border under both the cut and window interpretations, denoted by *F*_*C*_(*T, θ*) and *F*_*W*_ (*T, θ*), respectively.

When treating the border as a cut, we label the neurons left of *F*_*C*_(*T, θ*) that reside in *F*_*W*_ (*T, θ*) as members of class, *K*_1_ and analogously label the neurons on the right of *F*_*C*_(*T, θ*) as *K*_2_. We can generate these labels for many (*T, θ*) pairs (i.e. construct many fake cuts). A classifier that is fit on such a labeling performs best when neurons in *K*_1_ and neurons in *K*_2_ look functionally and structurally disparate. Thus, if the border can best be characterized as a cut, we would expect the classifier to perform best when fit on *F*_*C*_(0, 0).

When treating the border as a window, we label the neurons within *F*_*W*_ (*T, θ*) as 1 and outside as 0. Analogously, we can generate these labels for many (*T, θ*) pairs (i.e. construct many fake windows). We aim to see if the true border (i.e. *T* = 0, *θ* = 0) classifier performs better than its fake border counterparts. A classifier that is fit on such a labeling performs best when neurons within and outside the window look functionally and structurally disparate. Thus, if the border can best be characterized as a window, we would expect the classifier to perform best when fit on *F*_*W*_ (0, 0). We explore both interpretations of the border to determine which is most robust. See Appendix E for more details.

We explore two types of classifiers for both interpretations: a white box classifier and black box classifier. The white box classifier refers to manually constructing a feature representation of each window using some subset of structural and functional metrics discussed in Section 3.4. This form of classification is more interpretable, though limited in its expressivity. The black box classifier refers to utilizing a feature representation of each cut/window via less interpretable methods, as described below. Such a classifier enables us to motivate the border more thoroughly than the white-box classifier by capturing features we are unable to identify via the chosen set of structural and functional metrics.

To generate a white box structural feature representation of a given neuron *n*, we take a circular window around *n* with a radius of 20, 000 *µm* and compute a subset of the structural metrics outlined in Section 3.4. We can analogously generate a white box functional feature representation for a given neuron *n* by using a subset of the functional metrics outlined in Section 3.4.

To generate a black box structural representation of a given neuron *n*, we embed the graph using a variational-graph auto encoder (Kipf and Welling, 2016). A variational-graph auto encoder (VGAE) is designed to learn latent representations of neurons with higher order information in the graph that would be difficult to identify manually. Since the VGAE model only supports undirected graphs, we make the edges in the V1-RL graph bidirectional. Following training of the VGAE, a two dimensional vector encoding of each neuron is generated, which we use as the structural representation of *n*.

To generate a black box functional representation of a given neuron *n*, we use UMAP (McInnes et al., 2018) to reduce the dimensionality of the activity traces, as each trace contains 40, 000 measurements. While we lose interpretability along the temporal dimension, this approach has proven to be effective for classification modeling on functional data (Kumar et al., 2021). Finally, for the white-box models, we fit a logistic regression, and for the black-box models, we fit a multi-layer feed-forward neural network trained by gradient descent.

## 4. Results

### 4.1. Sliding window analysis reveals disparities between border regions

We use the sliding window framework (Section 3.5) to compare the structural and functional profiles of the V1-RL and RL-AL networks (Figure 5A), particularly at their respective borders. In terms of structure, we compute neuron density and synaptic density, and in terms of function, we find the correlation among the neural traces. For each sliding statistic that we compute, we normalize them to be between 0 and 1 using the minimum and maximum values. We do so separately across the two windows. This is to account for the potential differences in structure and function between the larger V1, RL and AL visual areas that we explore later (Section 4.2). Namely, we focus on the rate at which structure and function change across each network, particularly at the border region.

**Figure 5:**
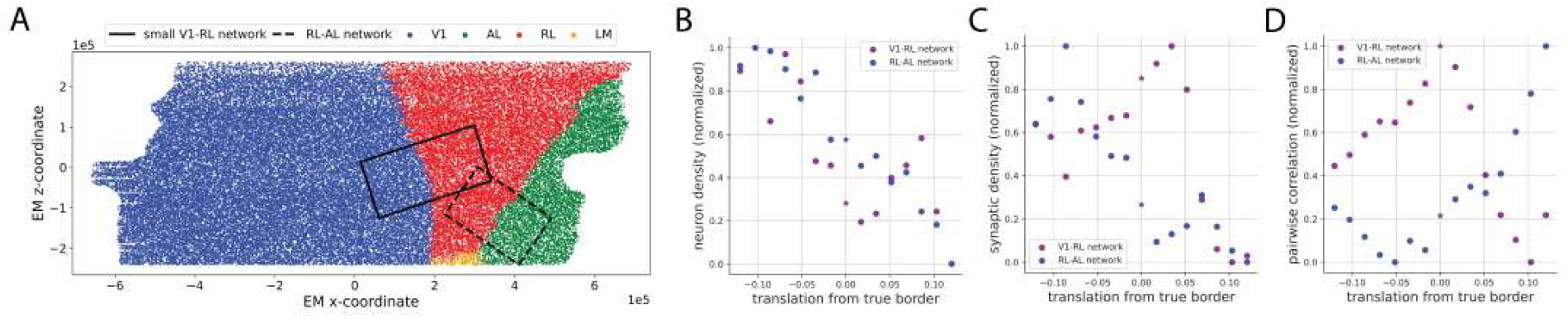
(A) Visualization of windows used for the comparison analysis relative to the full MICrONS graph. The solid black rectangle bounds the V1-RL window considered for this analysis. The dashed black rectangle bounds the RL-AL window used for this analysis. (B, C, D) Note that the values in the plots shown in (B), (C) and (D) are min-max normalized to explain away differences between the V1-RL and RL-AL networks. The star in each plot denotes the border region between the two visual areas. The x-axis refers to the distance of the sliding window away from the true border in *µm*. (B) Neuron density sliding window analysis. (C) Synaptic density sliding window analysis. (D) Pairwise correlation of functional activations measured in sliding window analysis.

**Figure 6:**
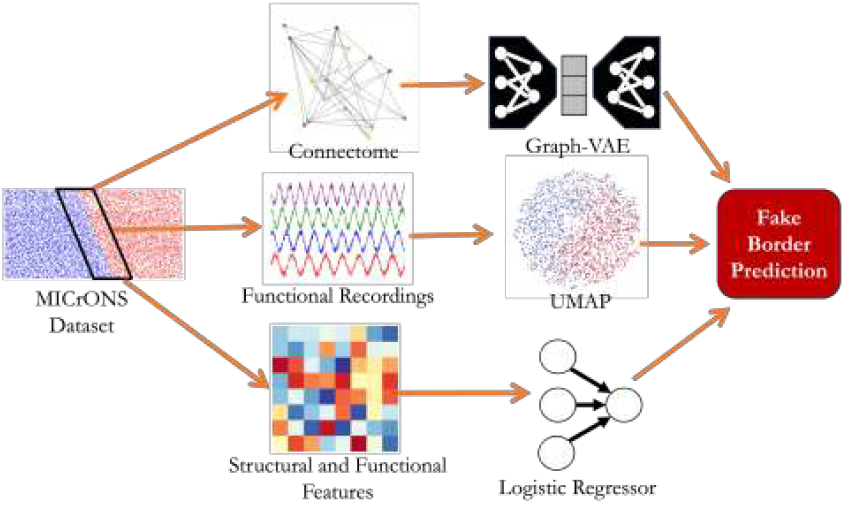
We extract the anatomical information and functional recordings of all neurons within a particular window and use both white-box and black-box classifiers to perform neuron-level classification across a set of fake borders. Here, we use graph variational auto encoders and UMAP as the inputs to black-box classifiers and logistic regression on manually selected features as a white-box classifier.

With respect to neuron density, we find similar trends across both networks in that neuron density decreases as we move from V1 to RL and RL to AL (Figure 5B). Interestingly, there appears to be a local minimum at the V1-RL border that we do not observe at the RL-AL border. When analyzing synaptic density, we note a more obvious difference between the two networks. The V1-RL border is a region of increased synaptic density, while the RL-AL border is not (Figure 5C). We leave further investigation of these differences to future work.

To compare the functional profile of the two networks, we analyze average pairwise correlation between the activity of neurons in the RL-AL and V1-RL regions. We find that the activity at the RL-AL border is not particularly notable compared to other sub-regions of the RL-AL graph (Figure 5D). In contrast, the V1-RL border is the most synchronous within the local V1-RL network. This suggests a functional disparity between the overall V1-RL and RL-AL network profiles, specifically at their border regions.

Overall, we have observed indications that the RL-AL border does not tend to present itself uniquely within the RL-AL network concerning structure or function (Figure 5). In contrast, the V1-RL border does present itself uniquely with respect both synaptic structure and function, perhaps acting as a standalone region in the local V1-RL network. Given the distinctiveness of the V1-RL border and the increased data availability of the V1 and RL regions, the remainder of this work will focus on analyzing the V1 and RL visual areas in more detail. We leave further exploration as to why different borders in the visual cortex are structurally and functionally distinct to future work.

### 4.2. Large scale differences between V1 and RL

In Section 4.1, we identify the V1-RL network and its border as a region of interest within the visual cortex. Here, we first aim to understand the V1 and RL regions as a whole beyond the metrics explored in Section 4.1. In particular, we investigate the metrics formalized in Section 3.4. For this and future analyses, since we are considering only the V1-RL network and are no longer constrained by the lack of data in the AL region, we expand our window of analysis (Figure 2).

In Section 4.1, we find that neuron density decreased from V1 to RL in the smaller portion of the V1-RL network (Figure 5A). We observe the same trend when analyzing the V1 and RL regions as a whole—neurons are more densely packed in V1 than in RL (Figure 7B). This is in line with previous, albeit less granular, observations in the primate visual cortex (Collins et al., 2010).

**Figure 7:**
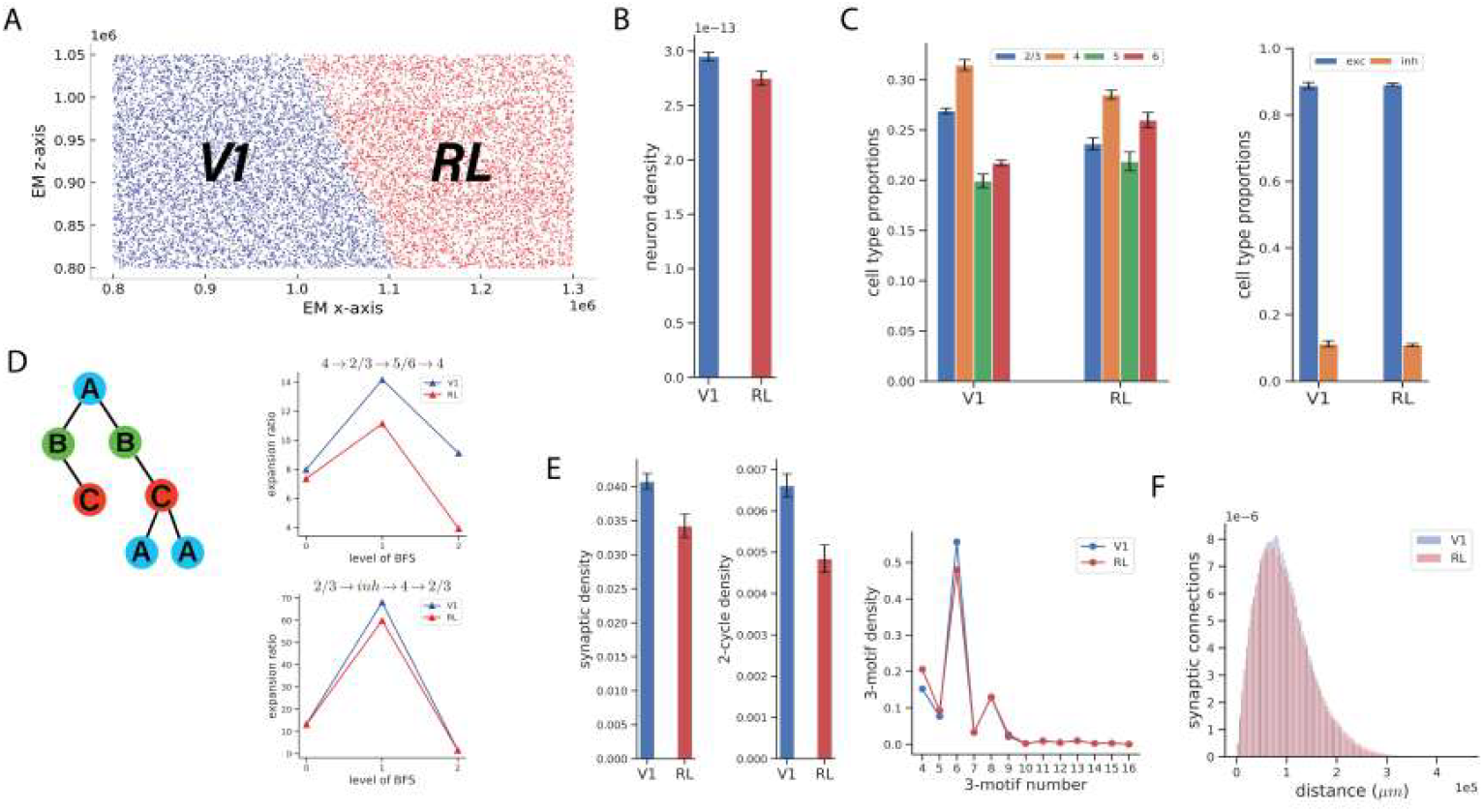
(A) Depiction of the V1-RL region we analyze. We estimate structural and function statistics over the entirety of V1 and RL. This is the same as the window shown in Figure 2, but plotted without the surrounding network. (B) Regional neuron density measured in units of 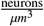. Two-sample t-test reveals higher neuron density in V1 (p-value *<* 0.01). (C, left) Neuron composition amongst excitatory types. Chi-squared goodness of fit test reveals difference between regions (p-value *<* 0.01). (C, right) Relative frequencies of inhibitory and excitatory neurons. Two-sample t-test shows no statistical difference between regions (p-value *>* 0.01). (D, left) Diagram showing an arbitrary example of the BFS tree generated by a microcircuit expansion. (D, top right) Microcircuit expansion rate for the excitatory microcircuit. Two-sample t-test conducted on each level of BFS reveals faster expansion in V1 (at least one level of BFS has p-value *<* 0.01) (D, bottom right) Microcircuit expansion rate for inhibitory microcircuit. Two-sample t-test conducted on each level of BFS faster expansion in V1 (at least one level has p-value *<* 0.01). (E, left) Regional synaptic density measured in units of 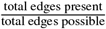. Two-sample t-test reveals higher synaptic density in V1 (p-value *<* 0.01). (E, middle) 2-cycle density measured in units of 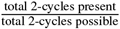. Two-sample t-test reveals higher 2-cycle density in V1 (p-value *<* 0.01). (E, right) Distribution over 3-motifs in each region. The 3-motif numbering is described in Section 3.4.5. Chi-squared goodness of fit test shows difference between regions (p-value *<* 0.01). For B, C, and E, we use bootstrapping as detailed in Section 3.4.1 with *t* = 0.2 and with 20 samples. (F) Distance-connectivity distributions for V1 and RL. Two-sample KS-test reveals longer-range connections in RL (p-value *<* 0.01).

In terms of cell type proportion differences, we find that L2/3 and L4 neurons occur more frequently in V1 and L5 and L6 in RL (Figure 7C, left). This aligns with the overall connectivity between different regions involved in visual processing. As V1 is the first cortical area to receive visual inputs from the thalamus, V1 may require more L4 neurons to process the inputs accurately. An analogous line of reasoning can be used for L2/3 neurons, which are also mediators of visual input.

Synaptic density is also higher in V1 than RL (Figure 7E, left), in line with the trend observed in the sliding window analysis conducted on the smaller V1-RL network (Figure 5B). We provide further insight on connectivity patterns by analyzing the relationship between the distance of any given pair of neurons and the probability a connection exists between them, finding that V1 tends to form shorter-range connections on average than RL (Figure 7F).

We move beyond basic measures of structure and examine 2-cycle and 3-motif properties of the network. In line with elevated synaptic density, we find that the density of reccurent 2-cycles is also higher in V1 (Figure 7E, middle). Another common measure of connectivity is 3-motifs: specific patterns of connectivity involving three nodes in a graph (Figure A1). These motifs have important implications for the functioning and behavior of biological networks. Some motifs might create paths for efficient information propagation, while others can lead to information bottlenecks or localized processing. While the specific functionality of each motif is still being refined, we find differences in the distributions of 3-motifs, specifically motifs 4 and 6. In particular, the former is more prevalent in RL, while the latter is more prevalent in V1 (Figure 7E, right). With its fan-out structure, motif 4 may indicate parallel and higher-order processing, while motif 6, with its chain structure, suggests a more ordered, hierarchical processing approach (Figure A1). This difference may indicate differences in information flow between the two regions, a thesis that we hope to explore in future work.

The final structural metric we analyze is microcircuit expansion. We find that the expansion rate of the microcircuits we investigated tends to be higher on average in V1 (Figure 7D). Taken together, our structural findings suggest that signals are propagated faster in V1 than in RL.

We now analyze the V1 and RL regions from a functional perspective, using the co-registered subset of neurons. We find that neuronal activity is higher in V1 than RL, as measured by the proportion of time a neuron fires, averaged across all neurons in the region (Figure 8B). Furthermore, we investigate the effective dimensionality of neural activity across the two regions. We find that the activity in RL is inherently lower dimensional than the activity in V1 (Figure 8C). Finally, we also observe a difference in the average pairwise correlation, aligning with the trends observed in the smaller V1-RL network analyzed previously (Figure 5D). In particular, we find that activity is more correlated in V1 (Figure 8D).

**Figure 8:**
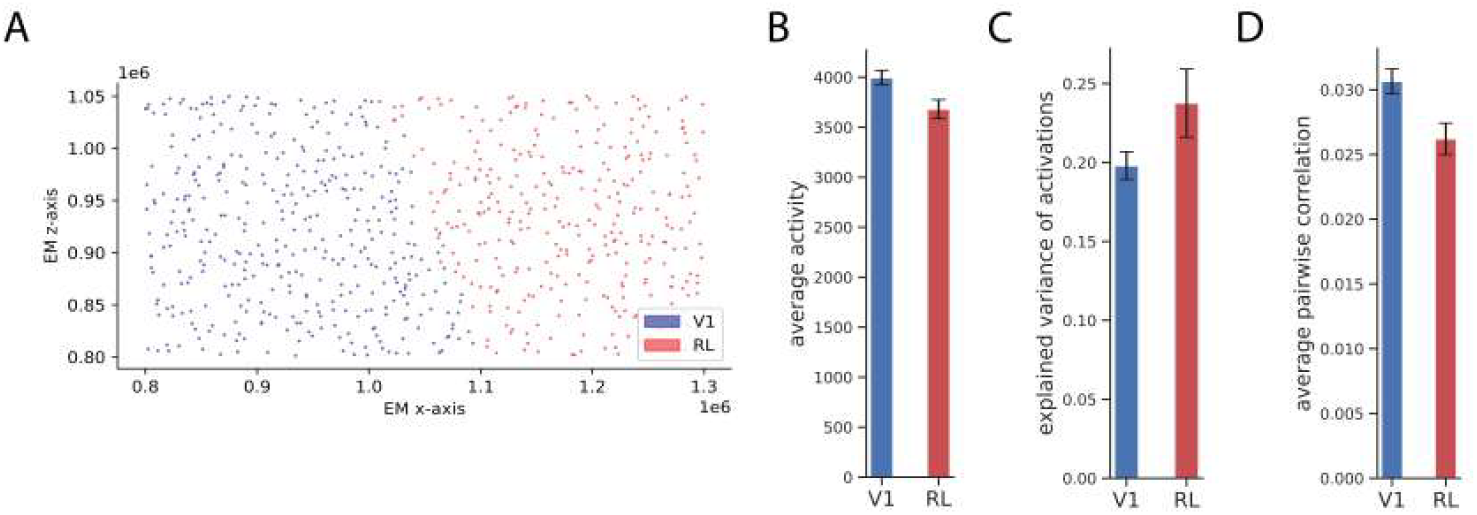
(A) Depiction of the co-registered dataset. Note how much sparser this is than the full graph in Figure 7A. All functional metrics are computed on this subgraph. (B) Average activity in V1 and RL. Two-sample t-test reveals higher activity level in V1 than RL (p-value *<* 0.01). (C) Effective dimensionality of activity in V1 and RL. Two-sample t-test reveals more complex activity in V1 than RL (p-value *<* 0.01). (D) Average pairwise correlation in V1 and RL. Two-sample t-test reveals more synchronous neuron firings in V1 than RL (p-value *<* 0.01). For B, C, and D, we use bootstrapping as detailed in Section 3.4.1 with *t* = 0.2 and with 20 samples.

Altogether, we speculate that these results arise from the notion of visual attention (Bisely, 2010). Visual attention selectively focuses on specific visual stimuli while filtering out others. The higher activity in V1 can reflect the prioritization of the brain to process the natural movie stimuli exposed to the mouse in greater detail. Due to the role of RL as a higher-order processor, it is unlikely that it necessitates the same computational capacity as V1, requiring both less activity and effective dimensionality. As a higher-order processing region, neurons within RL may abstract high-dimensional incoming data into lower-dimensional representations. This intuition is similar to what has been observed in artificial neural networks, specifically convolutional architectures. In such networks, earlier layers are tasked with extracting fine-grained details from the input image, and later layers filter these high-dimensional representations to generate a more robust, lower-dimensional representation useful for prediction (Recanatesi et al., 2019).

### 4.3. Granular analysis at interface of V1 and RL reveals border region

Next, we extend the sliding window framework to the larger portion of the V1-RL network and again analyze the change in various structural and functional metrics across the V1-RL region. Note that the shape of the sliding window is always constructed such that the edges are parallel to the border (Figure 9A), which here yields a different shape than was used in the analysis of the small portion of the V1-RL network (Section 4.1). This orientation is used to preserve as much data as possible (Figure 2, Section 3.1).

**Figure 9:**
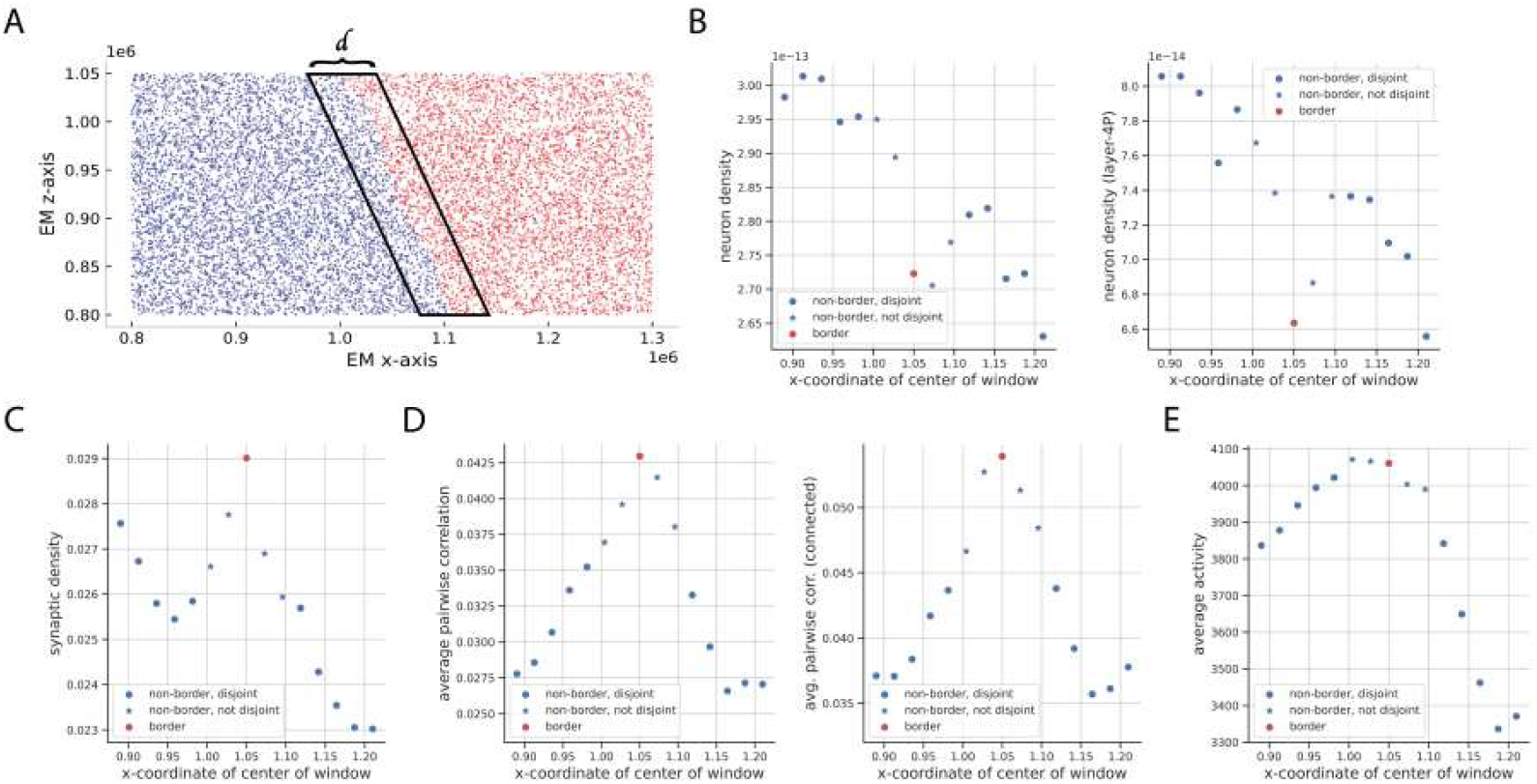
(A) Depiction of sliding window used in analyses that follow. (B, C, D, E) The legend in subsequent plots can be interpreted as follows: non-border, disjoint refers to windows that are disjoint from the border window and non-border, non-disjoint refers to windows that are not disjoint from the border window. Hence, the starred points represent a hybrid of the non-border and border subgraph. (B, left) Sliding window analysis for neuron density. (B, right) Sliding window analysis for L4 neuron density. We see that L4 neuron density decreases even more than the overall neuron density at the border. It is the only excitatory neuron type that decreases in this fashion at the border. (C) Sliding window analysis performed for synaptic density. (D, left) Sliding window analysis performed for average pairwise correlation computed amongst all pairs of neurons. (D, right) Sliding window analysis performed for average pairwise correlation only amongst connected neurons. (E) Sliding window analysis for average activity.

We again find that synaptic density presents itself uniquely at the V1-RL border, where it has the highest density among all sliding windows in the network (Figure 9C), offering support for our characterization of the V1-RL border as a more connected region of the visual cortex.

We analogously analyze neuron density and observe that at and around the border, the neuron density decreases significantly (Figure 9B, left), a trend even more pronounced than what we observed in Section 4.1. To understand this further, we analyzed the neuron density of each neuron type and found that the decrease at the border was most observable among L4 neurons (Figure 9B, right). Interestingly, the inverse relationship between synaptic density and neuron density occurs only at and around the border—synaptic density and neuron density are positively correlated in each retinotopic region at large and at other granular regions farther away from the border. This phenomenon warrants further investigation, and we currently hypothesize that it may partially be attributed to the border’s role as a message passer in the V1-RL network (we elaborate on this in Section 4.4). Since L4 neurons are sparse near the border, the firings may be dominated by information passed internally through the network (i.e. from V1 and RL) as opposed to externally from the thalamus.

Next, we examine functional metrics. Regarding average activity, we also observe a peak at the border (Figure 9E), similar to synaptic density. Previously, we observed a high-level relationship between connectivity and activity when aggregated over the entirety of V1 and RL (Section 4.2). Here, we observe a more granular relationship, where both synaptic density and average activity are highest at the border and lowest at the rightmost point in RL. Furthermore, we find that the average pairwise correlation between all neurons is highest at the border (Figure 9D, left), which is also in line with the same analysis performed on the smaller V1-RL network (Section 4.1). Similar to average activity, the change in pairwise correlation resembles the change in synaptic density over the network. Furthermore, when we consider correlation only between connected neurons, we find that the trend of the border being the region of highest pairwise correlation remains the same (Figure 9D, right).

To demonstrate the robustness of our characterization of the V1-RL border as a region distinct from V1 and RL, we conduct significance testing by spatially bootstrapping the V1, V1-RL border, and RL regions as described in Section 3.4. We find, both structurally and functionally, that the V1-RL border is statistically distinct from V1 and RL. Refer to Appendix C for more details.

### 4.4. Motifs in the V1-RL border region

Here, we attempt to develop intuition as to why the border is both structurally and functionally unique relative to the rest of the V1-RL network. We aim to identify various structural and functional motifs that arise at the border as a means of explaining its role in the visual cortex. The first motif we investigate captures the extent to which neurons communicate across the border cut (Figure 10A, left). This motif, which we term the cross-cut motif, is computed by measuring the Granger causality (a measure of information flow) of neurons connected across the cut for the various cuts constructed using the sliding window framework. We find that the true border contains the highest across-cut Granger causality (Figure 10A, right). The prevalence of the cross-cut motif at the border region may allude to its relevance as a conduit of information flow between V1 and RL.

**Figure 10:**
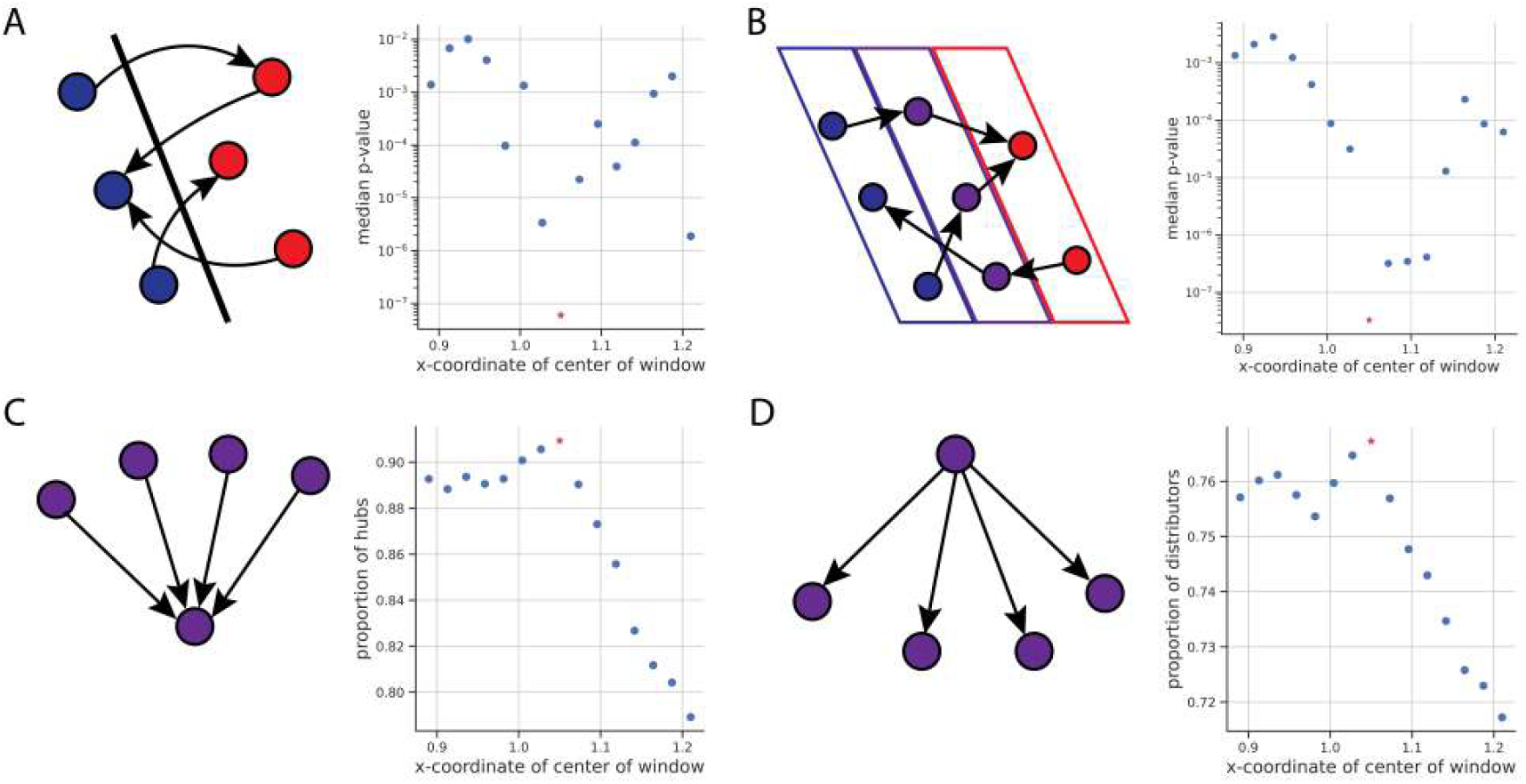
(A, left) Depiction of cross-cut motif. Median G-causality across all synapses crossing the cut is measured (using p-value as a proxy). (A, right) Cross-cut motif shows that information flow along synapses crossing the true border cut is the highest in the network. (B, left) Depiction of message passing motif. Median G-causality going from the blue neuron to the red neuron or from the red neuron to the blue neuron is measured depending on direction of connections. (B, right) We find that the message-passing motif is also highest at the border, indicating the role neurons in the border region play in carrying information between V1 and RL. (C, left) Depiction of hub motif (there being 4 pre-synaptic neurons in diagram is arbitrary). (C, right) The proportion of neurons that are hubs is highest near the border, pointing to the border’s role as an information aggregator. (D, left) Depiction of distributor motif (there being 4 post-synaptic neurons in diagram is arbitrary). (D, right) The proportion of neurons that are distributors is highest near the border, pointing to the border’s role as an information disseminator.

We further investigate this concept of message passing at the border by next looking at a message passing motif, which is a slight variant of the cross-cut motif. This motif attempts to show how well neurons preserve information flow from one to another through a third neuron, which is 3-motif number 6 (Figure A1). If three neurons are connected in a chain, *A* → *B* → *C*, then this motif measures the Granger causality between neurons *A* and *C* (Figure 10B, left). Neuron *B* can be viewed as a message passer between non-border neurons. Our analysis shows that message passing is significantly higher at the border than other areas in the network (Figure 10B, right).

Next, we analyze a related yet distinct set of border motifs: hubs and distributors. In the context of a biological network, a hub motif refers to a specific pattern or subgraph structure that represents a central node (the hub) connected to several pre-synaptic nodes (Figure 10C, left). These are related to 3-motifs 4 and 5 (Figure A1). Here, we interpret this motif as a unit of information aggregation. For our purposes, we arbitrarily define a hub motif to be a neuron that has 25 pre-synaptic neighbors. Similarly, a distributor motif refers to a central node (the distributor) connected to several post-synaptic nodes (Figure 10D, left). Here, we interpret this motif as a unit of information dissemination. We arbitrarily define a distributor motif to be a neuron that has 6 post-synaptic neighbors. We first examine the proportion of neurons that are hubs under the sliding window framework. In doing so, we find that hubs occur most frequently at the border region (Figure 10C, right). This supports the notion of the border acting as a region that is responsible for aggregating information from multiple different sources, and also offers a reason as to why the border is a highly connected subgraph of the V1-RL network. In addition, we also find that distributors occur most frequently at the border region (Figure 10D, right) which lends itself to the idea of the border acting not only as a region of information aggregation, but also information dissemination.

Overall, we now have a clearer picture of how the boundary between V1 and RL presents itself uniquely with respect to both structure and function. The cross-cut and message passing motifs both act as proxies for the strength of information flow and are highest at the border. Furthermore, the notion of message passing is associated with the ability of the border to aggregate and disperse information. By analyzing the hub and distributor motifs, we demonstrated that, structurally, the border region has the highest frequency of neurons that act to both absorb information from many sources and spread it to many destinations.

### 4.5. Characterization of V1-RL border as a standalone region is robust

Finally, we construct structural and functional white and black box classifiers for the V1-RL border region to demonstrate that our characterization of the border region as a standalone window is robust under perturbation. As described in Section 3, for the white box classifier, we generate a two-feature representation of each neuron that contains a measure of local synaptic and neuron density. We fit a classifier to each permutation of the labeling and find that the classifier is most accurate when distinguishing the real border from other neurons (Figure 11B). We then leverage neuron embeddings generated by a VGAE to train a classifier. Using the more complex representation shows that the classifier also best discerns the real border (Figure 11C).

**Figure 11:**
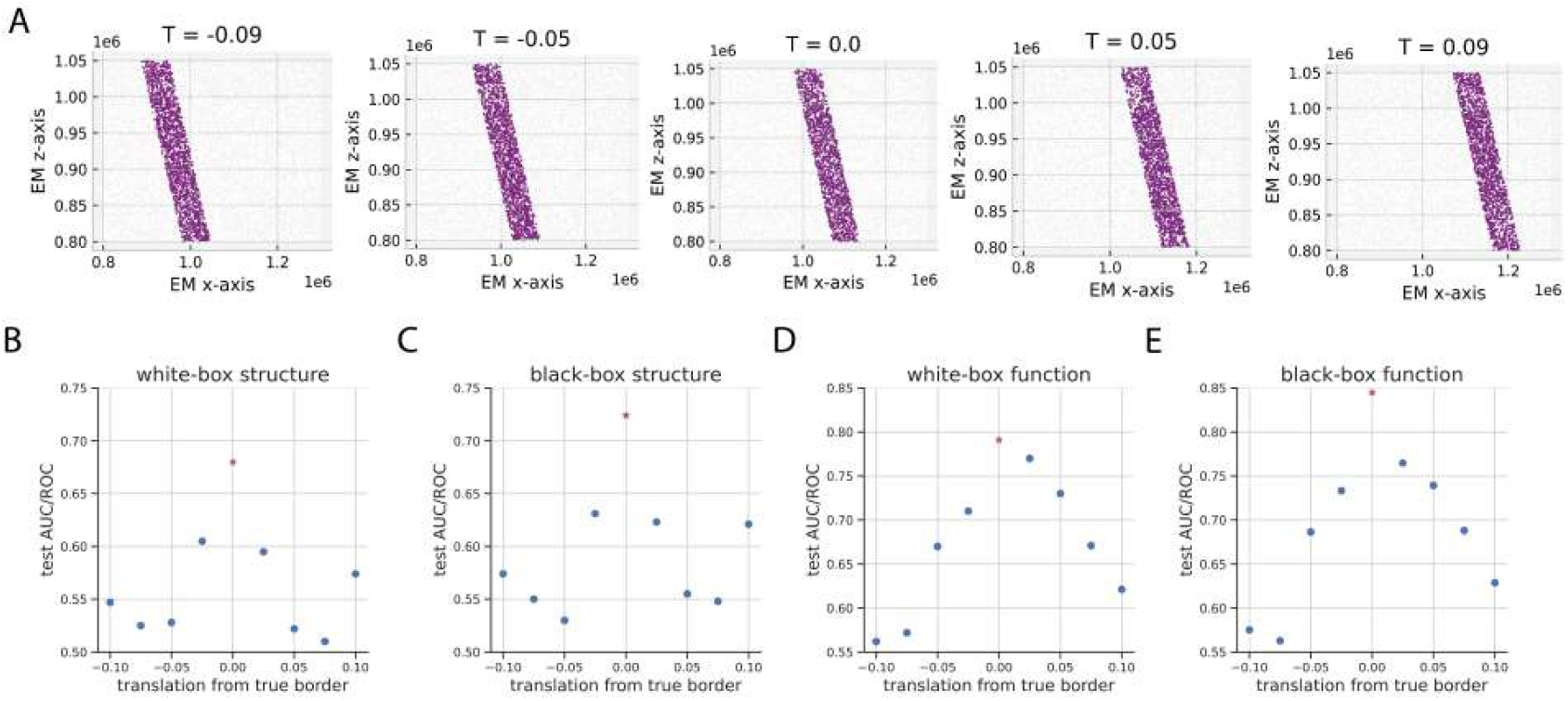
(A) Depiction of arbitrary fake borders constructed via horizontal translation. To perform classification for a given translation *T*, we label neurons within the purple window with a 1 and within other windows with a 0. The red dot in plots that follow denotes the true border. (B) Classifiers fit on a two-feature representation of each neuron using synaptic and neuron density. (C) Classifiers fit on a representation of neuron learned by VGAE. (D) Classifiers fit on a single-feature representation of each neuron using average pairwise correlation. (E) Classifiers fit on a neuron representation generated via UMAP.

We analogously construct a functional white box and black box classifier. For the white box classifier, we consider a single-feature representation of a neuron using the local average pairwise correlation. We also find that under this simple functional featurization, the border is identifiable (Figure 11D). We then construct a more complex functional featurization via UMAP embeddings of the activity traces of each neuron, and also find that the classifer performs best on the real border (Figure 11E). We also conducted additional experiments, which demonstrate that the real border region is also most identifiable under rotational perturbations (refer to Appendix D for these results). These results suggest the robustness of the V1-RL border region as a distinct subregion within the V1-RL network. Across both manually selected and model generated features, we find that the true border region yields the greatest classification accuracy.

Previously, we discussed an analogous form of fake border analysis under the cut interpretation, where the border acts as a 1-D decision boundary between V1 and RL (Section 3.3). However, throughout this work, we have adopted the window interpretation of the V1-RL border, where the border acts as 2D standalone region, without evaluating the feasibility of window interpretation relative to the cut interpretation. To validate our proposed window interpretation against the more canonical cut interpretation, we conduct the fake border analysis described in Section 3.6, among others. The results and discussion for this comparison analysis can be found in Appendix E.

## 5. Related Work

Due to recent advancements in automated neuronal segmentation, more and more comprehensive connectomics datasets are being released, such as those of the worm (Witvliet et al., 2021), fly (Scheffer et al., 2020; Dorkenwald et al., 2024), zebrafish (Svara et al., 2022), mouse (Consortium et al., 2021; Turner et al., 2022), and human (Shapson-Coe et al., 2024). Many other analyses build upon these datasets, investigating network statistics (Lin et al., 2024), inhibitory circuit patterns (Schneider-Mizell et al., 2023), and functional microcircuits (Matheson et al., 2022). However, previous datasets with both connectivity and functional data coupled together are small in scale, or they lack functional data altogether. MICrONS presents the first—and at the time of this writing the only—dataset with neuronal connectivity and activity coregistered at such a large scale, which is why we focus our analysis on this dataset.

Additionally, the availability of connectomes has facilitated much research on artificial neural networks with empirically derived connectivity data but simulated neuronal behavior (Ding et al., 2024; Lappalainen et al., 2024; Wang et al., 2024). Even before connectomes were as available, previous works utilized statistical priors to generate informed connectivity diagrams and used such wirings for their simulation work (Markram et al., 2015; Shi et al., 2022). In this work, we limit our analysis to non-generated, empirical data to investigate existing properties of the boundaries between visual regions. However, in future work, we plan to create and utilize artificial models of neuronal behavior to better understand the functional implications of our findings.

Finally, our use of the MICrONS dataset to analyze the different visual regions broadly falls into previous retinotopic and architectonic bodies of work. As introduced earlier, one standard approach to defining visual areas in the mouse visual cortex is using a retinotopic mapping (Bibollet-Bahena et al., 2023). Such maps give rise to borders between regions which are linked to changes in the chirality of the retinotopic map, identified functionally through the recording of neuronal responses using electrodes or imaging techniques (Zhuang et al., 2017). Another approach to mapping the visual cortex is architectonics, which unlike retinotopy is instead concerned with the structural organization of the brain, providing information about the arrangement of neurons, cell types, and cortical layers (Olavarria and Montero, 1989; Herculano-Houzel et al., 2013). Borders constructed using this approach are associated with rapid shifts in the aforementioned structural metrics. The presence of discussion on whether there is a mismatch between the boundaries of visual regions discerned through retinotopy and architectonics (Schuett et al., 2002; Zhuang et al., 2017) demonstrates the need for a more robust and nuanced formulation of structure and function in the visual cortex. sIn particular, retinotopic analyses primarily operate in a functional coordinate space (Kalatsky and Stryker, 2003), while architectonic ones operate in a physical coordinate space. Connectomics consolidates structural and functional data into a single, shared coordinate space via co-registration, enabling the analysis of structure and function in tandem. Additionally, retinotopy, which primarily focuses on localized responses to visual stimuli, may not inherently capture broader network dynamics and connectivity properties; this is of particular importance when analyzing boundaries between visual areas in the network. Finally, connectomics surpasses architectonics in providing a more detailed and comprehensive view of brain structure by capturing multiscale connectivity patterns and detailed circuitry.

Overall, our work attempts to leverage the more nuanced profiling of the visual cortex conferred by connectomics by understanding how we can use it to improve upon existing characterizations enabled by retinotopy and architectonics. While many previous works have connected structure and function (Chavane et al., 2011; Furutachi et al., 2023; Haber et al., 2023), ours is among the first to do so at the scale and resolution afforded by connectomics to analyze the boundaries between visual cortex regions.

## 6. Conclusion and Limitations

Understanding the mouse visual cortex, its retinotopically and architectonically derived visual regions, and the discrepancies between the two methodologies is an important step towards improving our understanding of the brain. In this work, we propose a statistical sliding window framework for analyzing neural networks in connectomics datasets like MICrONS that utilize both the connectivity graph and functional activations. By placing our focus on the borders between visual areas, our findings reveal two overarching themes: one, that not all borders are the same, and two, that the V1-RL border in particular serves as a distinct, possibly standalone piece of the visual cortex. For the latter theme, we find further characterization of the V1-RL border alludes to its role as a message passing interface within the larger V1-RL network.

Unfortunately, our work is constrained statistically in two primary ways. First, there are, to our knowledge, no other connectomics datasets publicly available at the scale of MICrONS that includes both structural and functional information. Therefore, we cannot make comparisons of our results on MICrONS with other datasets, limiting the generalizability of our findings. Second, even with the scale of MICrONS, due to our focus of the boundaries between visual cortex regions, our ability to perform significance testing of all of our results is hampered. Even so, we are able to perform some statistical significance testing comparing the V1 and RL regions to the border in terms of neuronal and synaptic density as well as functional activity (Section Appendix C).

In future work, we plan to make use of recently released foundation models for neuronal functional behavior (Wang et al., 2024). In this way, we can quickly test hypotheses that explore what functional implications the unique properties at the visual cortex boundaries allow for. Furthermore, we plan to develop a more extensive profiling of the visual cortex that includes analysis of its other regions and respective borders using our sliding window approach, beyond what is explored in Section 4.1. Since this relies upon a more extensive profiling of the visual cortex than is afforded by MICrONS in its current form, we hope to extend our work using later versions of the dataset as well as other datasets. We hope to extend our analytical pipeline to be more general than simply exploring boundaries as well. Finally, we hope to translate these results into artificial neural network design by inducing an interface for communication between subnetworks that functionally specialize, in order to emulate the role the V1-RL border region plays in the mouse visual cortex.

Overall, our use of the extensive MICrONS connectomics dataset sheds new light on what was previously a disparity between retinotopic and architectonic parcellations of the visual cortex. While there are limitations to our work, we hope that the analytical framework we introduce as well as the initial insights it yields inspires future exploration in this area.

## 7. Acknowledgments

We would like to thank Nancy Kanwisher, Bruce Fischl, and Jonathan Polimeni for their very helpful comments and discussions regarding our analysis. We would also like to thank Forrest Collman for his assistance with accessing and utilizing the MICrONS dataset. Finally, we would like to thank Lu Mi for her advice and guidance on the project.

This work was supported by an NIH Brains CONNECTS U01 grant.

## Appendix A. Neuron Types

The neurons in the dataset can be categorized as either excitatory or inhibitory. For the purposes of our analyses, we do not distinguish between subgroups amongst inhibitory neurons. Within the excitatory class of neurons, we consider only pyramidal neurons (abbreviated as P throughout the text) which belong to one of the following four cortical layers:

- **Layer 2*/*3 (L2*/*3) - External Granular Layer:** Contains small pyramidal neurons and stellate cells; plays a role in integrating sensory information.
- **Layer 4 (L4) - Internal Granular Layer:** Receives direct input from the thalamus and is critical for processing visual information from the eyes.
- **Layer 5 (L5) - External Pyramidal Layer:** Contains large pyramidal neurons that project locally and to other brain regions; involved in relaying information and motor control.
- **Layer 6 (L6) - Polymorphic Layer:** Contains a mix of neurons with diverse functions, sending feedback signals to the thalamus and other areas.

## Appendix B. Diagrams of all 3-motifs

**Figure A1:**
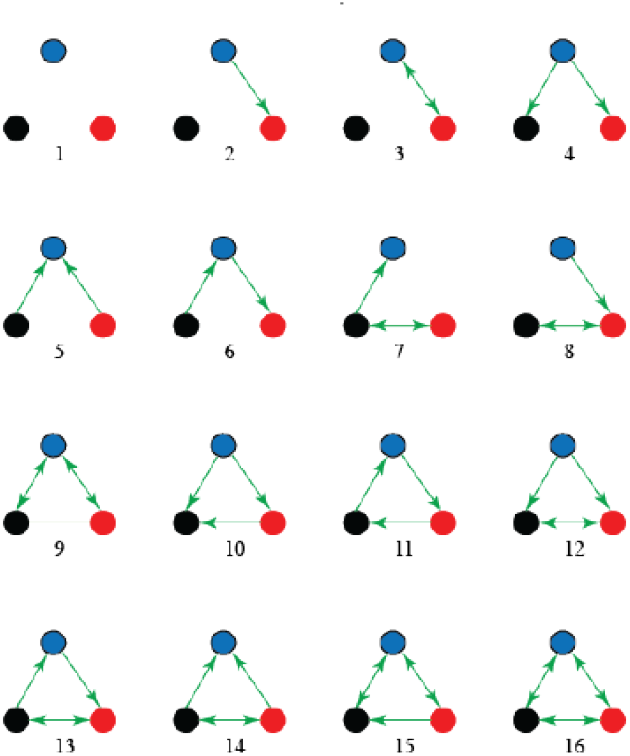
Set of all structural motifs formed by three neurons.

## Appendix C. Significance testing for V1-RL border

Since the windows in the sliding window analysis discussed in Section 4.3 do not contain much data to subsample, to demonstrate the statistical significance of the claim that the V1-RL border is structurally and functionally distinct from the V1 and RL regions, we divide the V1-RL network into these 3 regions (V1, RL, and the V1-RL border), and compute statistics on each of them.

For structure, we analyze synaptic density, neuron density and layer 4P neuron density. We do so by spatially bootstrapping the graph (as described in Section 3.4) and use the resulting set of data points to conduct 2 distinct two-sample t-tests: one comparing the V1-RL border to V1 and the other comparing the V1-RL border to RL.

Regarding function, we compute the average pairwise correlation between neural traces. Since we have trace data for many sessions (Consortium et al., 2021), we can instead use the individual sessions to construct multiple samples of pairwise correlation for the t-tests. Thus, we do not have to spatially bootstrap the graph and instead can consider the full regions. Again, here, we conduct 2 distinct two-sample t-tests: one comparing the V1-RL border to V1 and the other comparing the V1-RL border to RL.

We find that the V1-RL border is statistically distinct from both V1 and RL for all the aforementioned measures, except for overall neuron density (Figure A3). For neuron density, we find that while the V1-RL border is distinct from V1, it is not statistically distinct from RL. This is why we turn to measuring layer-4P neuron density as a structural measure to distinguish the border region from V1 and RL (Section 4.3).

**Figure A2:**
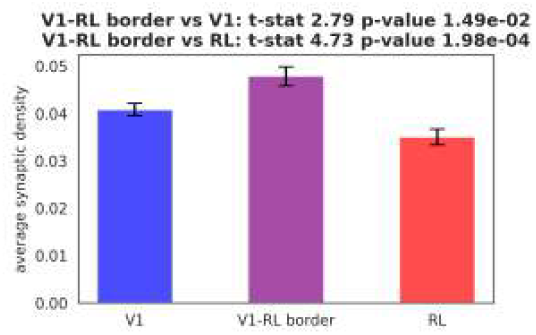
Synaptic connectivity statistical significance results from 2 sample t-test demonstrate that V1-RL border is more connected relative to V1 and RL.

**Figure A3:**
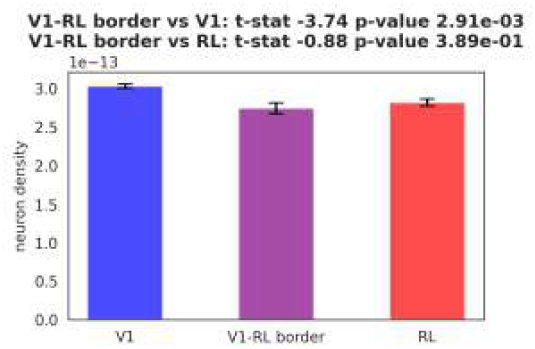
Overall neuron density statistical significance results demonstrate that V1-RL border is more neuronally sparse than V1. However, it is not statistically distinct from RL.

**Figure A4:**
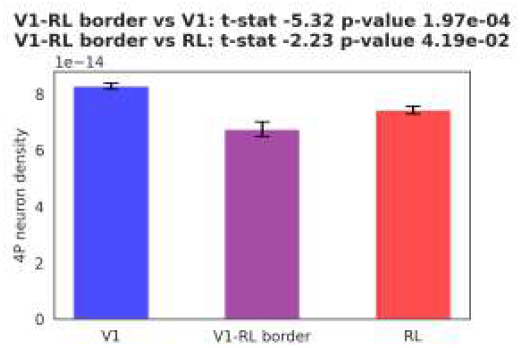
Layer-4P neuron density statistical significance results demonstrate that V1-RL border is more sparse than V1 and RL with respect to layer 4P neurons.

**Figure A5:**
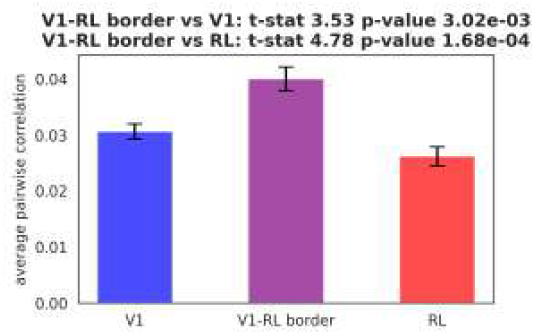
Average pairwise correlation analysis demonstrates that V1-RL border has significantly more synchronous neural activity than V1 and RL.

## Appendix D. Identifiability of V1-RL border under rotation

Recall that in Section 4.5, we showed that the true border is identifiable amongst fake borders constructed by horizontal translation. Here, we additionally show that the border is identifiable against fake borders constructed by rotating the true border around its center point. Before developing a classification framework that addresses this question, we will revisit the sliding window framework for analysis. Recall that the shape of the window is constructed such that the bounds of the window are parallel to the border axis. We can analogously construct sliding windows for fake borders generated by rotation and compute the sliding statistics under the newly shaped borders (Figure A6A).

With respect to structure, we compute both the sliding synaptic density and neuron density across rotations of the border window. We find that rotations of the border to the left (*θ <* 0) no longer have a distinct synaptic density at the border. However, for rotations to the right, there exist some rotated borders in which the fake border also presents itself uniquely with respect to synaptic density (Figure A6B). While one could argue that the synaptic density at the border window for rotation *θ* = 0 presents itself most uniquely across all generated rotations, it is not a strong argument for border identifiability. So, we next aim to construct a more robust set of identifiability criteria.

To do so, we turn to sliding neuron density across rotations of the border. Specifically, we look at L4 neuron density since we know that L4 density is responsible for the decrease in overall neuron density observed at the border (Figure 9B, left). Interestingly, we find that rotations of the border to the right (*θ <* 0) no longer have a distinct L4 neuron density at the border (Figure A6C). Recall that rotations of the border to the left no longer resemble the true border with respect to sliding neuron density. So, using both synaptic density and neuron density, the true border is identifiable against rotation. We formalize the notion of border identifiability against rotation by constructing a classifier that leverages these structural differences. Neurons are featurized by taking a neighborhood around them and computing the synaptic and L4 neuron density. For each rotation, we construct a classifier under the labeling *L*_0_ in which the true border is given a label of 1 and the rotated borders are given a label of 0. As expected, we find that the true border is more identifiable than its fake counterparts constructed by rotation (Figure A6F).

Next, we perform an analogous form of analysis using functional data. We first compute the sliding pairwise correlation across various rotations of the border window. The correlation at the true border appears to present itself most uniquely relative to the fake border rotations, however there exist a few rotations for which pairwise correlation is also highest at the border (Figure A6D). So, we analyze a potentially more robust metric in Granger causality (Section 3.4). Granger causality moves beyond pairwise correlation by capturing the temporal dynamics of network activity. Again, we perform a sliding window analysis using the fake rotated borders and find there is less ambiguity with respect to the identifiability of the true border than we observed analyzing pairwise correlation (Figure A6E). To demonstrate the identifiability of the true border using functional data, we featurize neurons by computing their local neighborhood pairwise correlation and Granger causality (as discussed in Section 3) and perform classification under the same paradigm discussed for the structural rotation classifier. In doing so, we find that the true border is more identifiable than the rotated fake borders (Figure A6G).

**Figure A6:**
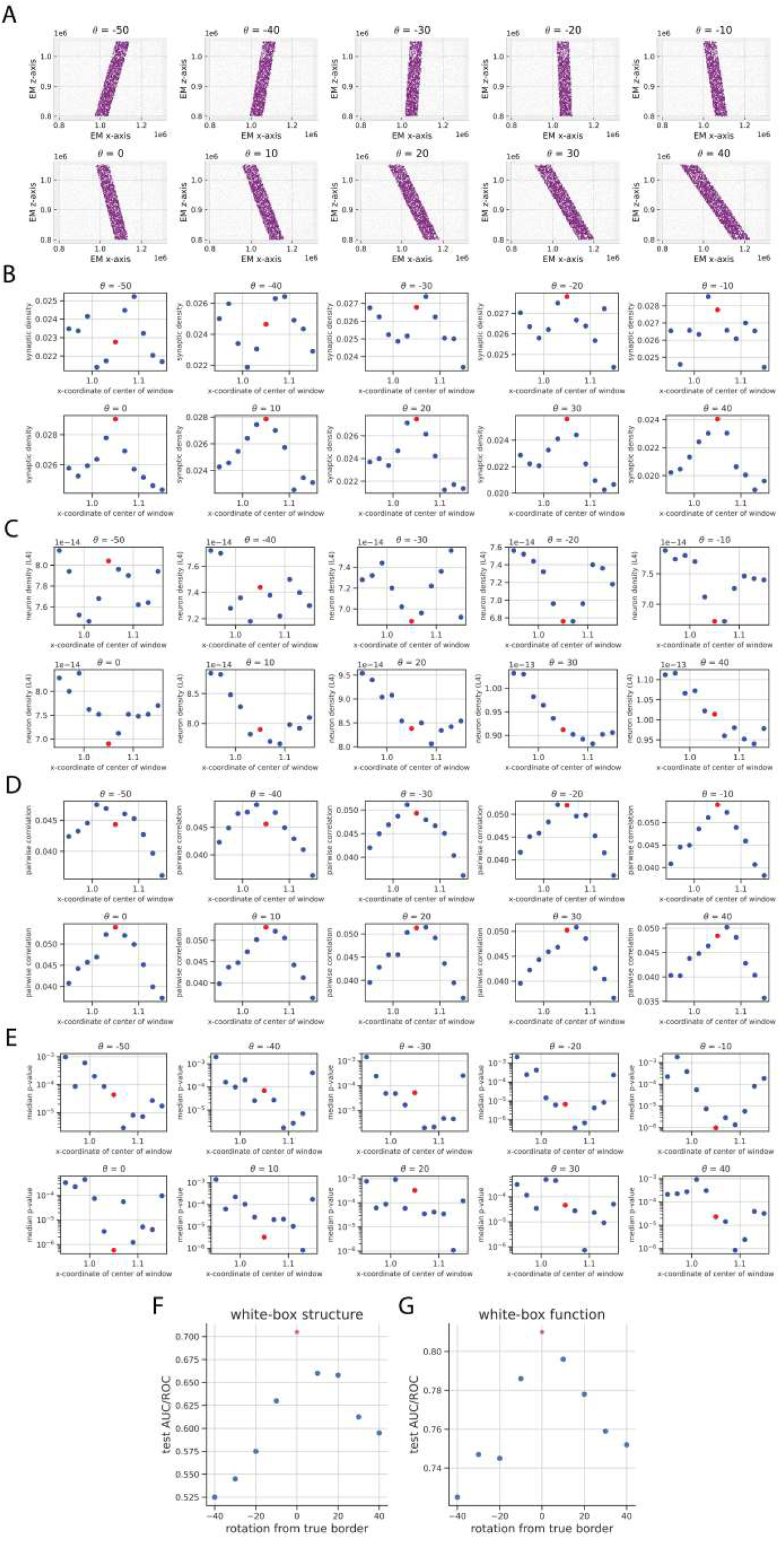
(A) Depiction of arbitrary fake rotations of the true border. We use these rotations in both the sliding window and classification frameworks discussed in previous analyses. Red dot denotes true border in all the following plots. (B) Sliding window analysis measuring synaptic density across various border rotations. (C) Sliding window analysis measuring L4 neuron density. (D) Sliding window analysis measuring average pairwise correlation amongst connected neurons. (E) Sliding window analysis measuring median Granger causality (using p-value as a proxy). (F) Structural notable magnitude classifier fit across multiple rotations of border. (G) Functional notable magnitude classifier fit across multiple rotations of border.

## Appendix E. Comparing the cut and window interpretations

Throughout the main text (particularly in Section 3.3), we have referenced this notion of interpreting the border as a cut and as a window. In this section, we make this notion more rigorous by providing analysis that supports the window interpretation over the cut interpretation. Furthermore, we offer more granular potential characterizations of border regions between visual areas and delineate the set of possible interpretations for the role they can play within their respective networks.

### Appendix E.1. Contextualizing the cut and window interpretations

Here, we provide some additional context to our characterization of the cut and window interpretations of the V1-RL border. Recall in Section 3.3 we described the cut interpretation as a decision boundary between V1 and RL whereas the window interpretation characterizes the V1-RL border as a standalone region in the visual cortex. To make this idea more precise, we consider two paradigms and frame each as a classification problem (as described in Section 3.6): the border as a region of notable rate of shift and as a region of notable magnitude.

By notable rate of shift, we mean that with respect to some structural or functional measure, the border denotes a region in the V1-RL network over which the rate at which this measure changes is high relative to the rest of the network. This follows naturally from how the V1-RL border is currently fit via retinotopic parcellation. Namely, since a parcellation constitutes a classification problem with respect to the retinotopic labels, we can interpret the fitted border as a decision boundary between the V1 and RL networks. We will refer to the border as a region of notable rate of shift and the border under the cut interpretation interchangeably.

By notable magnitude, we mean that with respect to some structural or functional measure, the border denotes a region in the V1-RL network over which this measure is high or low relative to the rest of the network. Note that this characterization of the border follows less readily from the canonical fitting of the V1-RL border via a retinotopic parcellation. Since parcellation is synonymous with finding a decision boundary, the fitting procedure is aligned with the cut interpretation and fails to take into account the possibility that the V1-RL border itself may act as a distinct region within the larger V1-RL network. We will refer to the border as a region of notable magnitude and the border under the window interpretation interchangeably.

In Section 3.6, we discussed experimental schemes to examine the border under these two paradigms. In Sections 4.5 and Appendix D, we demonstrated that under the notable magnitude (i.e. window interpretation) characterization of the border, the true V1-RL border is identifiable amongst fake borders. Furthermore, the statistical analysis presented in Sections 4.3 and 4.4 showed that the border window presents itself uniquely with respect to a variety of structural and functional measures. Overall, we have essentially assumed the window interpretation with our sliding window approach and bolstered the analyses we have conducted. However, we now go one step further and conduct an analogous fake border analysis under the cut interpretation to demonstrate quantifiably why the window interpretation of the V1-RL border is more reasonable than the cut interpretation.

#### Appendix E.2. True V1-RL border is unidentifiable when interpreted as a region of notable shift

Here, we conduct a fake cut analysis analogous to the fake window analysis conducted in Section 4.5 and demonstrate that, in contrast to the window interpretation, under the cut interpretation, the border is not identifiable.

Recall that the rate of shift characterization of the border is synonymous to the cut interpretation of the border which lends itself most readily to existing interpretations of borders under retinotopic maps. In this regime, the border window serves simply as a decision boundary between V1 and RL. Since these two regions are functionally and structurally distinct (Section 4.2), we would expect the border to denote the region in which structure and function change the most. Yet, we know from earlier analyses (Section 4.3) that this interpretation of the border is unlikely, as both structural and functional statistics measured at the border do not present themselves as a hybrid of V1 and RL.

To demonstrate the inefficacy of interpreting the border as a region of notable shift, we construct fake cuts by horizontal translation (Figure A7A). We consider the classification problem detailed in Section 3.6 in which we use the same label for neurons that lie on the same side of the cut. The structural white box classifier employs a two-feature representation of each neuron by computing the synaptic density and neuron density of a subgraph constructed by taking a fixed size neighborhood around each point. Under this featurization, we find that the classifier performs worse at the true border relative to many of the fake borders (Figure A7B). If we were to use performance under this classifier as a means of identifying the true border, we would conclude that many of the fake windows are more reasonably construed as the true border than the true border itself. This result is unsurprising in lieu of the sliding window neuron and synaptic density findings discussed in Section 4.3. However, it is still possible that there exists a structural statistic that we have not computed in which the border appears identifiable under this classification paradigm. To address this, we construct a black box classifier in which the neuron representations in the graph are learned by a VGAE which is capable of capturing more nuanced aspects of network structure. Even here, we find that the true border is unidentifiable (Figure A7C).

**Figure A7:**
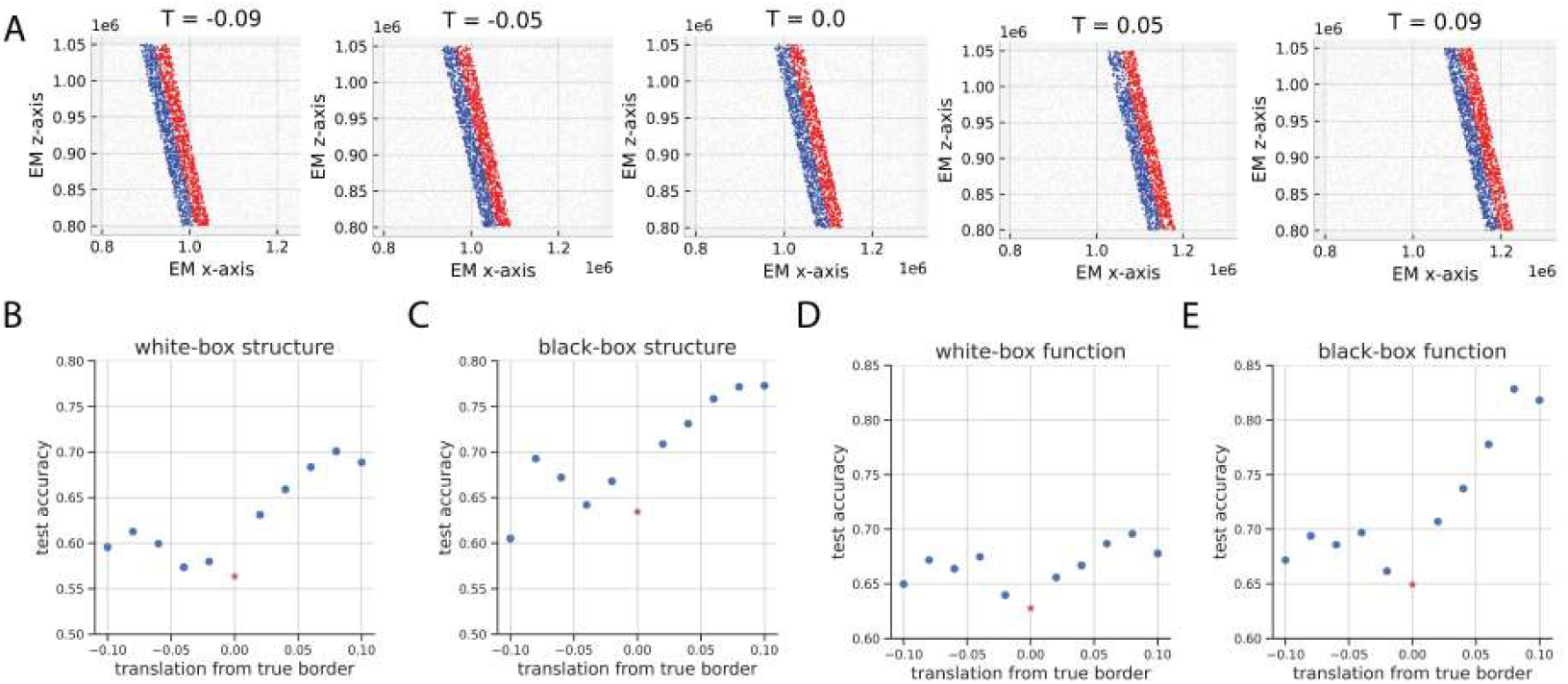
(A) Depiction of arbitrary fake border windows constructed via horizontal translation. To perform classification for a given translation, we label the neurons in blue with a 1 and the neuron in red with a 0. Recall that we refer to this classification paradigm as notable rate of shift. The red dot in plots that follow denotes the true border. (B) Classifiers fit on a two-feature representation of each neuron using synaptic and neuron density. (C) Classifiers fit on representation of neuron learned by VGAE. (D) Classifiers fit on a single-feature representation of each neuron using average pairwise correlation. (E) Classifiers fit on neuron representations generated via UMAP.

Next, we aim to see if the true border is also unidentifiable when using a functional feature representation of each neuron. Again, we construct fake cuts by horizontal translation. We generate a single-feature representation of each neuron by measuring the pairwise correlation of neurons in a subgraph constructed by taking a fixed size neighborhood around each point. Even under a functional feature representation, we find that the classifier does not perform particularly well at the true border (Figure A7D). Analogous to the structural white box classifier, this result is unsurprising given the sliding window pairwise correlation analysis (Figure 9). But, here we take a step further, and also show that a more expressive functional featurization is equally ineffective at identifying the border. Using UMAP on the neural traces yields better performance at fake borders than the true one (Figure A7E).

Recall that prior to this classification analysis, we knew that the notable rate of shift interpretation of the border was unlikely to be true given the observations we made in the sliding window analysis. However, here we made this notion concrete by showing that even when using more complex and expressive representations of neurons, we are still incapable of identifying the V1-RL border.

## Appendix F. Constructing more granular interpretations of the border

### Appendix F.1. Border regimes

**Figure A8:**
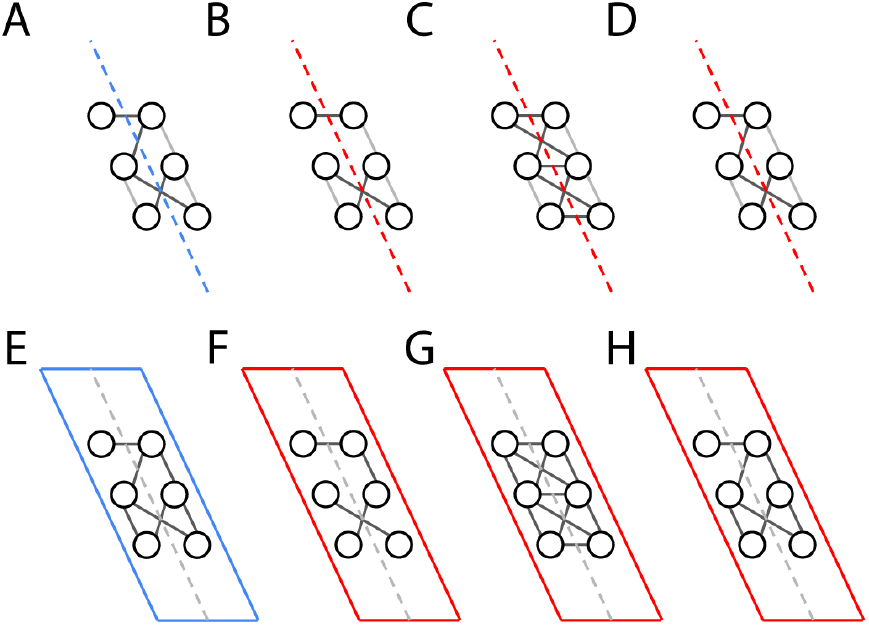
(A) Depiction of an arbitrary cut in the graph. Connections in black denote across cut connections, and connections in gray denote same side connections. We represent and discuss the other cut interpretations relative to this one. (B) Segregated cut regime. (C) Connected cut regime. (D) Homogenous cut regime. (E) Depiction of an arbitrary window in the graph. We represent and discuss the other window interpretations relative to this one. (F) Segregated window regime. (G) Connected window regime. (H) Homogenous window regime.

In this section, we propose more granular cut and window interpretations of the border, opening up a potential avenue for future work on characterizing the V1-RL border. Additionally, in Appendix F.3, we perform some analysis to contextualize the proposed interpretations, but note that our analysis is considered preliminary in nature.

In both the cut and window interpretations, we describe three possible regimes that potentially characterize the border purely in units of interactions between pairs of neurons: the segregated regime, the connected regime and the homogenous regime. We note that these interpretations can be generalized beyond interactions between just 2 neurons, but for the sake of simplicity, we restrict our scope as such. We first introduce these three regimes in the context of the cut interpretation.

1. **Segregated cut regime** Pairs of neurons interacting across the border cut are more structurally segregated and functionally dissimilar than the average pair of neurons sampled across an arbitrary cut in the graph (Figure A8B)
2. **Connected cut regime** Pairs of neurons interacting across the border cut are more structurally connected and functionally similar than the average pair of neurons sampled across an arbitrary cut in the graph (Figure A8C)
3. **Homogeneous cut regime** Pairs of neurons interacting across the border cut look structurally and/or functionally the same as the average pair of neurons sampled across an arbitrary cut in the graph (Figure A8D)

Analogously, we can introduce three possible regimes the state of the network can lie in under the window interpretation:

1. **Segregated window regime** Pairs of neurons interacting within the border window are more structurally segregated and functionally dissimilar than the average pair of neurons sampled from an arbitrary window in the graph (Figure A8F)
2. **Connected window regime** Pairs of neurons interacting within the border window are more structurally connected and functionally similar than the average pair of neurons sampled from an arbitrary window in the graph (Figure A8G)
3. **Homogeneous window regime** Pairs of neurons interacting within the border window look structurally and/or functionally the same as the average pair of neurons sampled from an arbitrary window in the graph (Figure A8H)

One thing to note is that in these interpretations, we implicitly assume structure and function are tied together.

#### Appendix F.2. Contrasting the border regimes

The window regimes proposed above encapsulate their respective cut regimes. In particular, the window regimes assign importance to all pairs of neurons in the window: this includes both pairs of neurons interacting across the cut and pairs of neurons interacting on the same side of the cut. In contrast, the cut regimes place emphasis only on the former.

In both the cut and window interpretations of the border, we have proposed three regimes for the state of a network. In the segregation regime, the border acts as a plane (cut interpretation) or region (window interpretation) over which information flow is significantly lower than the rest of the network. This is a natural hypothesis since it is reasonable to believe that two visual areas act individually as information processors and any information leakage from one area to the other would be detrimental to each regions’ ability to specialize as a functional unit.

In the connected regime, we posit that the border presents itself as a hub for information aggregation and dissemination – acting as a conduit of information flow between the two regions. This is also a reasonable hypothesis, as it suggests that while the two regions act as distinct information processors, their ability to communicate and synergize enhances computation in the visual cortex.

In the homogenous regime, the border does not present itself uniquely with respect to either structure or function, and its existence is simply induced by differences in retinotopy between the two regions. In this case, the border denotes the shift in structure and function when going from one region to another.

We note that amongst the three proposed border regimes, the canonical interpretation under existing formulations of a border between visual areas is the homogenous regime. In particular, the leading assumption in these formulations is that the border simply represents the region of greatest shift in structure and/or function in the network. In contrast, the segregated and connected regimes take a different stance: instead, they posit that the border itself is a distinct construct, potentially serving its own role within the larger network.

#### Appendix F.3. Structural and functional trends suggest V1-RL border lies in connected window regime

Up to this point, our results with respect to structural and functional properties have suggested that the window interpretation of the border is a more reasonable model than the cut interpretation (Section 4.3, 4.4). Here, we explicitly distinguish between neurons connected across the cut and on the same side of the cut (Figure A9D). We find that under synaptic density, the border is equally identifiable (Figure A9E) using cross-cut and same side connections. We notice the same effect when measuring pairwise correlation (Figure A9F). These findings reinforce the window interpretation of the border by demonstrating that the identifiability of the border is not solely attributable to structural and functional relationships formed across it—rather, the entire region around the border plane is unique. We hope to explore the idea that the V1-RL border lies in the connected window regime in future work.

#### Appendix F.4. V1-RL border as a localized region

Here, we provide further characterization of the V1-RL border lying in the connected border window regime by considering the notion of locality. In all of our previous analyses, we considered only connections between neurons contained within the window (Figure 8). An alternative formulation is to still consider neurons in the border window, but to allow them to form connections with neurons outside the window that are contained within a larger window (Figure A9A). We recompute both synaptic density and pairwise correlation under this non-local formulation and find that under both metrics, the distinct peak at the border we observe in the local formulation dissipates (Figures A9B, A9C). This suggests that the structural and functional uniqueness of the V1-RL border is a localized phenomenon. We hope to explore this notion of locality further in future work.

**Figure A9:**
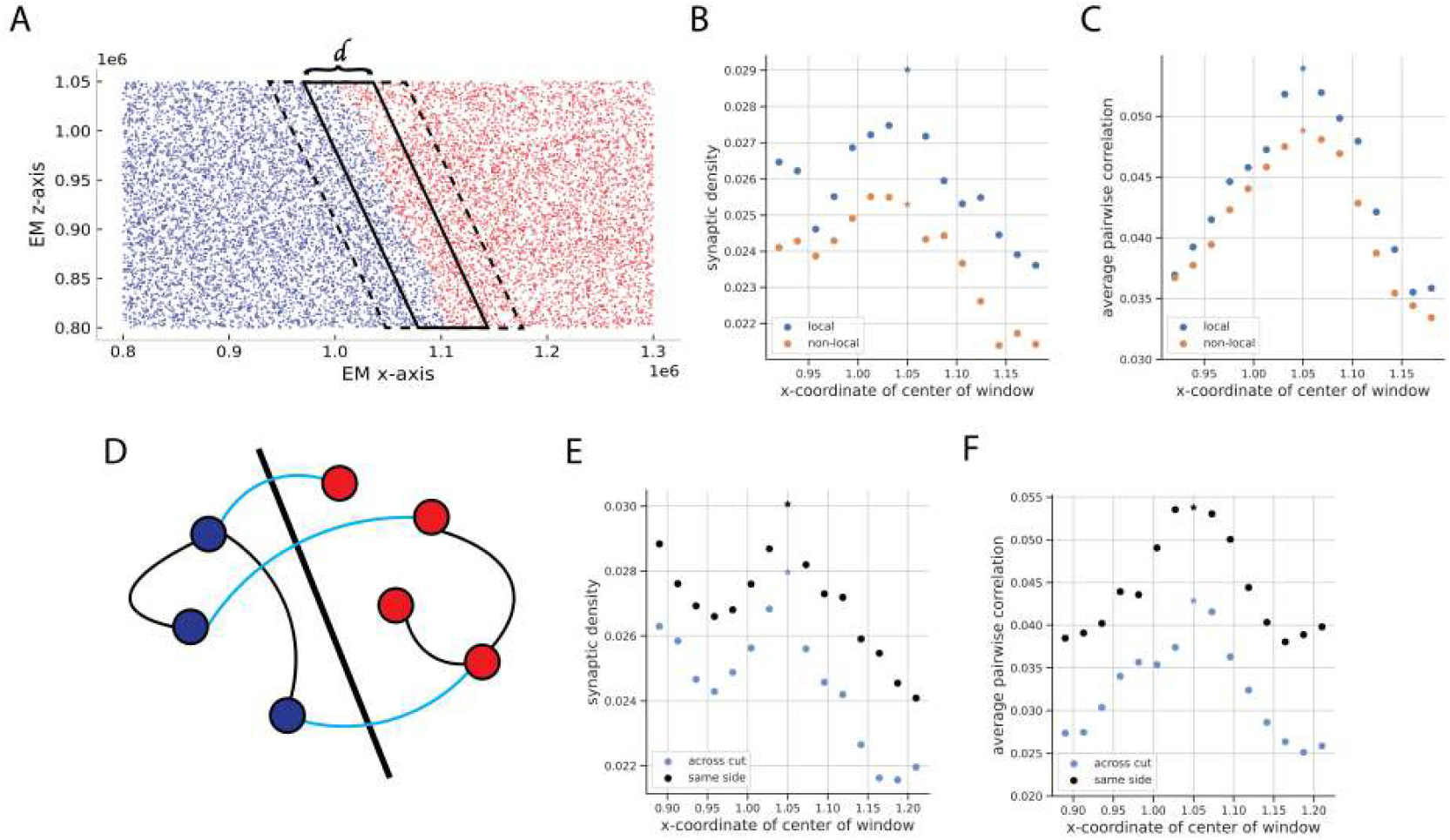
(A) Depiction of a secondary window that allows us to measure the locality/non-locality of the computed metrics. In this analysis, we examine all connections formed within the dashed-line window, such that at least one neuron lies in the solid-line window. In the plots that follow, the star denotes the border window. (B) Plot comparing local and non-local synaptic density. Local refers to sliding window analysis shown in Figure 9 in which only connections formed within the solid-line window are analyzed. (C) Locality analysis for average pairwise correlation. (D) Diagram showing the distinction between connections formed on the same side of the cut (black) and across the cut (light blue). (E) Comparison of the synaptic density of the subgraph formed by considering neurons across the cut versus the same side of the cut. (F) Comparison of the average pairwise correlation of the subgraph formed by considering neurons across the cut versus the same side of the cut.

## References

Baldauf, Z.B., 2005. Smi-32 parcellates the visual cortical areas of the marmoset. Neuroscience letters 383, 109–114.

Bastos, A.M., Usrey, W.M., Adams, R.A., Mangun, G.R., Fries, P., Friston, K.J., 2012. Canonical microcircuits for predictive coding. Neuron 76, 695– 711.

Behrens, T.E., Sporns, O., 2012. Human connectomics. Current opinion in neurobiology 22, 144–153.

Bibollet-Bahena, O., Tissier, S., Ho-Tran, S., Rojewski, A., Casanova, C., 2023. Enriched environment exposure during development positively impacts the structure and function of the visual cortex in mice. Sci Rep 13, 7020.

Bisely, J., 2010. The neural basis of visual at-tention - PubMed — pubmed.ncbi.nlm.nih.gov. https://pubmed.ncbi.nlm.nih.gov/20807786/. [Accessed 28-02-2025].

Chavane, F., Sharon, D., Jancke, D., Marre, O., Frégnac, Y., Grinvald, A., 2011. Lateral spread of orientation selectivity in v1 is controlled by intracortical cooperativity. Frontiers in systems neuroscience 5, 4.

Chen, X., Ginoux, F., Carbo-Tano, M., Mora, T., Walczak, A.M., Wyart, C., 2023. Granger causality analysis for calcium transients in neuronal networks, challenges and improvements. eLife 12, e81279. doi:10.7554/eLife.81279.

Collins, C.E., Airey, D.C., Young, N.A., Leitch, D.B., Kaas, J.H., 2010. Neuron densities vary across and within cortical areas in primates. Proc Natl Acad Sci U S A 107, 15927–15932.

Consortium, M., Bae, J.A., Baptiste, M., Bishop, C.A., Bodor, A.L., Brittain, D., Buchanan, J., Bumbarger, D.J., Castro, M.A., Celii, B., et al., 2021. Functional connectomics spanning multiple areas of mouse visual cortex. BioRxiv, 2021–07.

Ding, Z., Fahey, P.G., Papadopoulos, S., Wang, E.Y., Celii, B., Papadopoulos, C., Chang, A., Kunin, A.B., Tran, D., Fu, J., et al., 2024. Functional connectomics reveals general wiring rule in mouse visual cortex. bioRxiv, 2023–03.

Dorkenwald, S., Matsliah, A., Sterling, A.R., Schlegel, P., Yu, S.C., McKellar, C.E., Lin, A., Costa, M., Eichler, K., Yin, Y., Silversmith, W., Schneider-Mizell, C., Jordan, C.S., Brittain, D., Halageri, A., Kuehner, K., Ogedengbe, O., Morey, R., Gager, J., Kruk, K., Perlman, E., Yang, R., Deutsch, D., Bland, D., Sorek, M., Lu, R., Macrina, T., Lee, K., Bae, J.A., Mu, S., Nehoran, B., Mitchell, E., Popovych, S., Wu, J., Jia, Z., Castro, M., Kemnitz, N., Ih, D., Bates, A.S., Eckstein, N., Funke, J., Collman, F., Bock, D.D., Jefferis, G.S.X.E., Seung, H.S., Murthy, M., 2023. Neuronal wiring diagram of an adult brain. bioRxiv.

Dorkenwald, S., Matsliah, A., Sterling, A.R., Schlegel, P., Yu, S.C., McKellar, C.E., Lin, A., Costa, M., Eichler, K., Yin, Y., et al., 2024. Neuronal wiring diagram of an adult brain. Nature 634, 124–138.

Fize, D., Vanduffel, W., Nelissen, K., Denys, K., d’Hotel, C.C., Faugeras, O., Orban, G.A., 2003. The retinotopic organization of primate dorsal v4 and surrounding areas: a functional magnetic resonance imaging study in awake monkeys. Journal of neuroscience 23, 7395–7406.

Furutachi, S., Franklin, A.D., Mrsic-Flogel, T.D., Hofer, S.B., 2023. Co-operative thalamocortical circuit mechanism for sensory prediction errors. bioRxiv, 2023–07.

Garrett, M.E., Nauhaus, I., Marshel, J.H., Callaway, E.M., 2014. Topography and areal organization of mouse visual cortex. Journal of Neuroscience 34, 12587–12600.

Van der Gucht, E., Hof, P.R., Van Brussel, L., Burnat, K., Arckens, L., 2007. Neurofilament protein and neuronal activity markers define regional architectonic parcellation in the mouse visual cortex. Cerebral Cortex 17, 2805– 2819.

Haber, A., Wanner, A., Friedrich, R.W., Schneidman, E., 2023. The structure and function of neural connectomes are shaped by a small number of design principles. bioRxiv, 2023–03.

Herculano-Houzel, S., Watson, C., Paxinos, G., 2013. Distribution of neurons in functional areas of the mouse cerebral cortex reveals quantitatively different cortical zones. Frontiers in neuroanatomy 7, 35.

Hooks, B.M., Chen, C., 2020. Circuitry Underlying Experience-Dependent Plasticity in the Mouse Visual System. Neuron 107, 986–987.

Jazayeri, M., Ostojic, S., 2021. Interpreting neural computations by examining intrinsic and embedding dimensionality of neural activity. Curr Opin Neurobiol 70, 113–120.

Kalatsky, V.A., Stryker, M.P., 2003. New paradigm for optical imaging: temporally encoded maps of intrinsic signal. Neuron 38, 529–545.

Kipf, T.N., Welling, M., 2016. Variational graph auto-encoders. arXiv preprint 1611.07308.

Kumar, M.G., Hu, M., Ramanujan, A., Sur, M., Murthy, H.A., 2021. Functional parcellation of mouse visual cortex using statistical techniques reveals response-dependent clustering of cortical processing areas. PLOS Computational Biology 17, e1008548.

Lappalainen, J.K., Tschopp, F.D., Prakhya, S., McGill, M., Nern, A., Shinomiya, K., Takemura, S.y., Gruntman, E., Macke, J.H., Turaga, S.C., 2024. Connectome-constrained networks predict neural activity across the fly visual system. Nature 634, 1132–1140.

Lin, A., Yang, R., Dorkenwald, S., Matsliah, A., Sterling, A.R., Schlegel, P., Yu, S.c., McKellar, C.E., Costa, M., Eichler, K., et al., 2024. Network statistics of the whole-brain connectome of drosophila. Nature 634, 153– 165.

Lyon, D.C., Xu, X., Casagrande, V.A., Stefansic, J.D., Shima, D., Kaas, J.H., 2002. Optical imaging reveals retinotopic organization of dorsal v3 in new world owl monkeys. Proceedings of the National Academy of Sciences 99, 15735–15742.

Markram, H., Muller, E., Ramaswamy, S., Reimann, M.W., Abdellah, M., Sanchez, C.A., Ailamaki, A., Alonso-Nanclares, L., Antille, N., Arsever, S., et al., 2015. Reconstruction and simulation of neocortical microcircuitry. Cell 163, 456–492.

Marshel, J.H., Garrett, M.E., Nauhaus, I., Callaway, E.M., 2011. Functional specialization of seven mouse visual cortical areas. Neuron 72, 1040–1054.

Matheson, A.M., Lanz, A.J., Medina, A.M., Licata, A.M., Currier, T.A., Syed, M.H., Nagel, K.I., 2022. A neural circuit for wind-guided olfactory navigation. Nature communications 13, 4613.

McInnes, L., Healy, J., Saul, N., Großberger, L., 2018. Umap: Uniform manifold approximation and projection. Journal of Open Source Software 3, 861. doi:10.21105/joss.00861.

Nassi, J.J., Callaway, E.M., 2006. Multiple circuits relaying primate parallel visual pathways to the middle temporal area. Journal of Neuroscience 26, 12789–12798.

Olavarria, J., Montero, V.M., 1989. Organization of visual cortex in the mouse revealed by correlating callosal and striate-extrastriate connections. Vis Neurosci 3, 59–69.

Recanatesi, S., Farrell, M., Advani, M., Moore, T., Lajoie, G., Shea-Brown, E., 2019. Dimensionality compression and expansion in deep neural networks. URL: https://arxiv.org/abs/1906.00443, 1906.00443.

Scheffer, L.K., Xu, C.S., Januszewski, M., Lu, Z., Takemura, S.y., Hayworth, K.J., Huang, G.B., Shinomiya, K., Maitlin-Shepard, J., Berg, S., et al., 2020. A connectome and analysis of the adult drosophila central brain. elife 9, e57443.

Schneider-Mizell, C.M., Bodor, A., Brittain, D., Buchanan, J., Bumbarger, D.J., Elabbady, L., Kapner, D., Kinn, S., Mahalingam, G., Seshamani, S., et al., 2023. Cell-type-specific inhibitory circuitry from a connectomic census of mouse visual cortex. bioRxiv.

Scholtens, L.H., van den Heuvel, M.P., 2018. Multimodal Connectomics in Psychiatry: Bridging Scales From Micro to Macro. Biol Psychiatry Cogn Neurosci Neuroimaging 3, 767–776.

Schuett, S., Bonhoeffer, T., Hübener, M., 2002. Mapping retinotopic structure in mouse visual cortex with optical imaging. Journal of Neuroscience 22, 6549–6559.

Seth, A.K., Barrett, A.B., Barnett, L., 2015. Granger causality analysis in neuroscience and neuroimaging. J Neurosci 35, 3293–3297.

Shapson-Coe, A., Januszewski, M., Berger, D.R., Pope, A., Wu, Y., Blakely, T., Schalek, R.L., Li, P.H., Wang, S., Maitin-Shepard, J., et al., 2024. A petavoxel fragment of human cerebral cortex reconstructed at nanoscale resolution. Science 384, eadk4858.

Shi, J., Tripp, B., Shea-Brown, E., Mihalas, S., A. Buice, M., 2022. Mousenet: A biologically constrained convolutional neural network model for the mouse visual cortex. PLOS Computational Biology 18, e1010427.

Svara, F., Förster, D., Kubo, F., Januszewski, M., Dal Maschio, M., Schubert, P.J., Kornfeld, J., Wanner, A.A., Laurell, E., Denk, W., et al., 2022. Automated synapse-level reconstruction of neural circuits in the larval zebrafish brain. Nature Methods 19, 1357–1366.

Thompson, M.E., Ramirez Ramirez, L.L., Lyubchich, V., Gel, Y.R., 2016. Using the bootstrap for statistical inference on random graphs. Canadian Journal of Statistics 44, 3–24. doi:10.1002/cjs.11271.

Turner, N.L., Macrina, T., Bae, J.A., Yang, R., Wilson, A.M., Schneider-Mizell, C., Lee, K., Lu, R., Wu, J., Bodor, A.L., et al., 2022. Reconstruction of neocortex: Organelles, compartments, cells, circuits, and activity. Cell 185, 1082–1100.

Tyler, C.W., 2004. Representation of stereoscopic structure in human and monkey cortex. Trends in Neurosciences 27, 116–118. doi:10.1016/j.tins.2003.12.009.

Tyler, C.W., Likova, L.T., Chen, C.C., Kontsevich, L.L., Schira, M.M., Wade, A.R., 2005. Extended concepts of occipital retinotopy. Current Medical Imaging 1, 319–329.

Wang, E.Y., Fahey, P.G., Ding, Z., Papadopoulos, S., Ponder, K., Weis, M.A., Chang, A., Muhammad, T., Patel, S., Ding, Z., et al., 2024. Foundation model of neural activity predicts response to new stimulus types and anatomy. bioRxiv, 2023–03.

Wang, Q., Gao, E., Burkhalter, A., 2007. In vivo transcranial imaging of connections in mouse visual cortex. Journal of neuroscience methods 159, 268– 276.

Witvliet, D., Mulcahy, B., Mitchell, J.K., Meirovitch, Y., Berger, D.R., Wu, Y., Liu, Y., Koh, W.X., Parvathala, R., Holmyard, D., et al., 2021. Connectomes across development reveal principles of brain maturation. Nature 596, 257– 261.

Zhuang, J., Ng, L., Williams, D., Valley, M., Li, Y., Garrett, M., Waters, J., 2017. An extended retinotopic map of mouse cortex. eLife 6, e18372. doi:10.7554/eLife.18372.

Zilles, K., Palomero-Gallagher, N., 2017. Multiple transmitter receptors in regions and layers of the human cerebral cortex. Frontiers in neuroanatomy 11, 78.

